# Interpretable Forecasting of Kidney Cancer Progression via Generative AI and Symbolic Reasoning

**DOI:** 10.64898/2026.08.23.746526

**Authors:** Guillermo Prol-Castelo, Elina Syrri, Nikolaos Manginas, Vasileios Manginas, Jon Sánchez-Valle, Nikos Katzouris, Georgios Paliouras, Alfonso Valencia, Davide Cirillo

## Abstract

Predicting cancer stage progression from omics data, and deriving molecular insight into the mechanisms driving it, remains a major challenge, owing in part to the lack of adequate longitudinal data and the interpretability limitations of current forecasting models. Large cancer datasets such as TCGA capture patient profiles cross-sectionally rather than longitudinally, complicating timely treatment decisions as tumors become more invasive. Deep neural networks typically used for forecasting, such as LSTMs, compound this problem by remaining largely opaque and offering clinicians no straightforward way to audit their predictions. Clear cell renal cell carcinoma (ccRCC) illustrates the clinical stakes of both challenges. Five-year survival falls from over 94% at stage I to 28% at stage IV, yet early-stage tumors are often managed under active surveillance, a strategy constrained by sparse molecular evidence of progression risk. Detecting progression in time, meanwhile, demands forecasts clinicians can interpret and trust, not black-box predictions. We address both challenges by combining generative and symbolic AI: a Variational Autoencoder trained on bulk RNA-Seq profiles of 530 TCGA ccRCC patients generates synthetic pseudotime trajectories that overcome the absence of longitudinal data, while a symbolic rule-induction framework (ASAL) learns finite-state automata from these trajectories, encoding stage transition as human-readable Boolean conditions over gene expression, which a complex event forecasting system (Wayeb) converts into probabilistic forecasts of stage advancement. An independent XGBoost classifier trained on real patients (F1 score = 0.71-0.81) shows a gradual early-to-late probability shift along the synthetic trajectories, absent in non-progressing control trajectories. Pathway enrichment of those trajectories reveals stage-dependent changes in established kidney cancer-related processes, including the TCA cycle and DNA repair. Finally, our symbolic forecaster nearly matches an LSTM baseline (macro F1 = 0.928 vs. 0.964), while additionally offering an inspectable rule set and a probability distribution over transition timing rather than a single opaque score. This work shows that generative and symbolic AI, paired together, can turn cross-sectional cohorts into a transparent, forecast-oriented framework for modeling disease progression, demonstrated here in ccRCC.

## 1 Introduction

Renal cell carcinoma (RCC) is the most common kidney malignancy in adults, with clear cell RCC (ccRCC) accounting for approximately 75% of cases [1]. Despite advances in targeted therapy and immunotherapy, prognosis remains strongly tied to disease stage at diagnosis: five-year survival rates fall from more than 94% for stage I to an average of 28% for stage IV [2]. Stage progression in ccRCC is accompanied by distinct shifts in gene expression programs, making bulk RNA sequencing a natural substrate for identifying molecular correlates of disease advancement. The Cancer Genome Atlas (TCGA) has provided a large, well-annotated bulk RNA-seq cohort for ccRCC (TCGA-KIRC), comprising over 500 patients with matched RNA-seq and clinical staging data, that enables population-level investigation of stage-associated gene expression [3]. However, transcriptomic association with stage does not imply mechanistic relevance. Statistical selection alone cannot distinguish genes that drive progression from those that merely correlate with it, yielding biomarker lists of uncertain biological and clinical meaning [4].

Numerous studies have derived prognostic and staging signatures from TCGA ccRCC transcriptomes using machine learning and statistical approaches [5–8]. While such methods can achieve strong discrimination performance, they share a fundamental limitation. They treat stage stratification as a static classification problem, comparing aggregate expression profiles between early and late stages without modeling the trajectory of progression between them. Identifying genes that forecast stage transitions, rather than merely distinguish established stages, requires a longitudinal perspective that bulk RNA-seq cohorts like TCGA cannot directly provide, as they capture single snapshots per patient. This calls for an interpretable framework that can reconstruct progression trajectories from cross-sectional data and extract the gene expression patterns that are most predictive of transition, rather than those that are simply differentially expressed at the endpoints.

Variational Autoencoders (VAEs) [9] have emerged as a powerful framework for learning compact representations of high-dimensional cancer transcriptomes [10], with demonstrated utility in tumor subtyping [11], survival analysis [12], pseudo-time ordering in single-cell data [13], and the study of stage differences in latent space [14, 15]. Crucially, the generative capacity of VAEs enables synthetic data generation, yet its potential to reconstruct longitudinal progression trajectories remains largely unexplored in cancer research [16]. Such trajectories can be built by generating synthetic intermediate patients that bridge the cross-sectional snapshots available in TCGA, using demographically-matched source-target pairings to isolate stage-related change and multiple plausible endpoints per patient to reflect the inherent uncertainty in disease progression. The synthetic trajectories, however, are only valuable if one can extract from them interpretable patterns that forecast stage transitions. Forecasting disease progression has been constrained by the scarcity of longitudinal genomic data [17], and most approaches rely on static classification [18–20] or black-box sequential models such as LSTMs [21, 22] that, while predictive, offer limited transparency and no natural way to incorporate biological domain knowledge [23]. Beyond raw predictive accuracy, clinical adoption of AI-based forecasting tools depends on their transparency and auditability. Post hoc explanations of black-box models are not guaranteed to faithfully reflect the model’s actual decision process, motivating calls for models that are inherently interpretable by design in high-stakes domains such as healthcare [24]. Calibrated confidence, a model’s ability to indicate not only a prediction but how much that prediction should be trusted, is similarly recognized as a prerequisite for safely deferring uncertain cases to clinician judgment [25], yet standard deep learning models are frequently poorly calibrated and overconfident even as their raw accuracy improves [26]. These considerations motivate our design choice to evaluate the forecasting stage of our pipeline not only on predictive accuracy, but on the transparency and probabilistic nature of its predictions, properties we argue are equally consequential for eventual clinical deployment. Hybrid neuro-symbolic models address this by combining deep learning with logic-based components [27], and have shown promise in tasks such as entity recognition in clinical records [28] and diabetes prediction [29]. Nevertheless, their application to temporal forecasting of disease progression remains scarce [30, 31], and none have leveraged generative trajectories as the substrate for symbolic pattern learning.

To our knowledge, this is the first framework to combine VAE-generated longitudinal trajectories with symbolic reasoning for cancer stage progression forecasting. A concurrent preprint [32] similarly explores neuro-symbolic cancer progression forecasting, but relies on real (not synthetic) longitudinal expression data and a different symbolic technique (Algorithmic Information Dynamics). Here, we present a pipeline, summarized in Figure 1, that unifies these two capabilities: a VAE trained on TCGA-KIRC data reconstructs longitudinal gene-expression trajectories by generating synthetic patients that capture the progression from early to late stages; symbolic finite-state automata (SFA) [33] are then inferred from these trajectories, encoding stage-transition logic as interpretable Boolean combinations of gene-expression predicates; finally, the learned automata are embedded within a Complex Event Forecasting framework [34] that issues probabilistic, time-to-event estimates of stage advancement from partial trajectory observations. The result is a biologically transparent pipeline that identifies gene expression patterns most predictive of ccRCC stage progression and provides clinically interpretable forecasts, recovering gene programs consistent with established ccRCC biology, moving beyond snapshot-based association toward a mechanistically grounded model of disease dynamics. Throughout, we treat transparency and the availability of interpretable, probabilistic forecasts as design goals on par with predictive accuracy, given their direct relevance to safe clinical deployment.

**Fig. 1:**
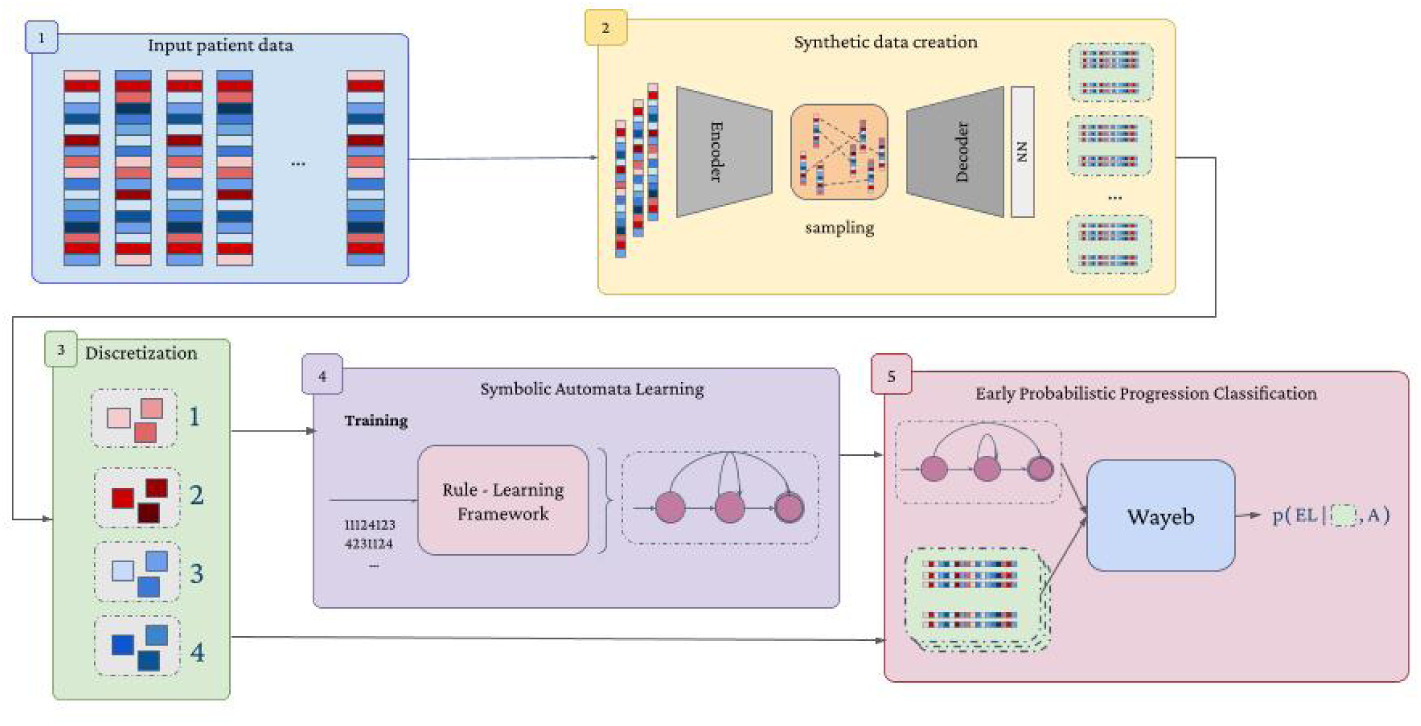
Synthetic trajectory modeling and interpretable forecasting of ccRCC stage progression. After preprocessing, the bulk-RNA-Seq data from the TCGA-KIRC dataset contains 530 ccRCC patients. This data is used to train a VAE with a post-processing neural network for synthetic data generation. The synthetic data represent pseudotemporal molecular trajectories, modeling the gene expression changes across the stage transition between early and late cancer stages. ASAL (Answer Set Automata Learning) then learns symbolic finite-state automata (SFA) from the trajectories, encoding gene-expression patterns as interpretable predicates that Wayeb, the Complex Event Forecasting (CEF) framework, uses for probabilistic stage-transition forecasting. The result is a clinically interpretable, rule-based forecast of ccRCC stage progression based on bulk RNA-seq data.

## 2 Results

### 2.1 Classifying ccRCC stages

We obtained bulk RNA-Seq and clinical staging data for all TCGA cancer types via the UCSC Xena browser (Methods 4.1), grouped stages I and II as early and III and IV as late, and classified early/late status for each of the top 10 most common histological types in TCGA (Supplementary Table S1). Each cancer type was preprocessed separately (Methods 4.1) and we applied an XGBoost classifier with 100 boosting rounds, optimized with Optuna for 100 trials, on a tenfold cross-validation. Different classification metrics showed ccRCC as the cancer with the clearest separation between stages (Supplementary Figure S2). Besides, ccRCC contains patients at all stages and relatively balanced compared to other cancers (Supplementary Table S2). A randomization experiment on the early and late stage patient labels confirmed the obtained classification results were higher than those expected by chance (Supplementary Figure S1). Further data preprocessing to remove outlier genes (Methods 4.1) yielded similar classification results (Figure 2a) on the panel of 8,516 genes used in the rest of this work. The best-performing XGBoost classifier was kept for downstream analyses (see ROC curve in Supplementary Figure S3).

**Fig. 2:**
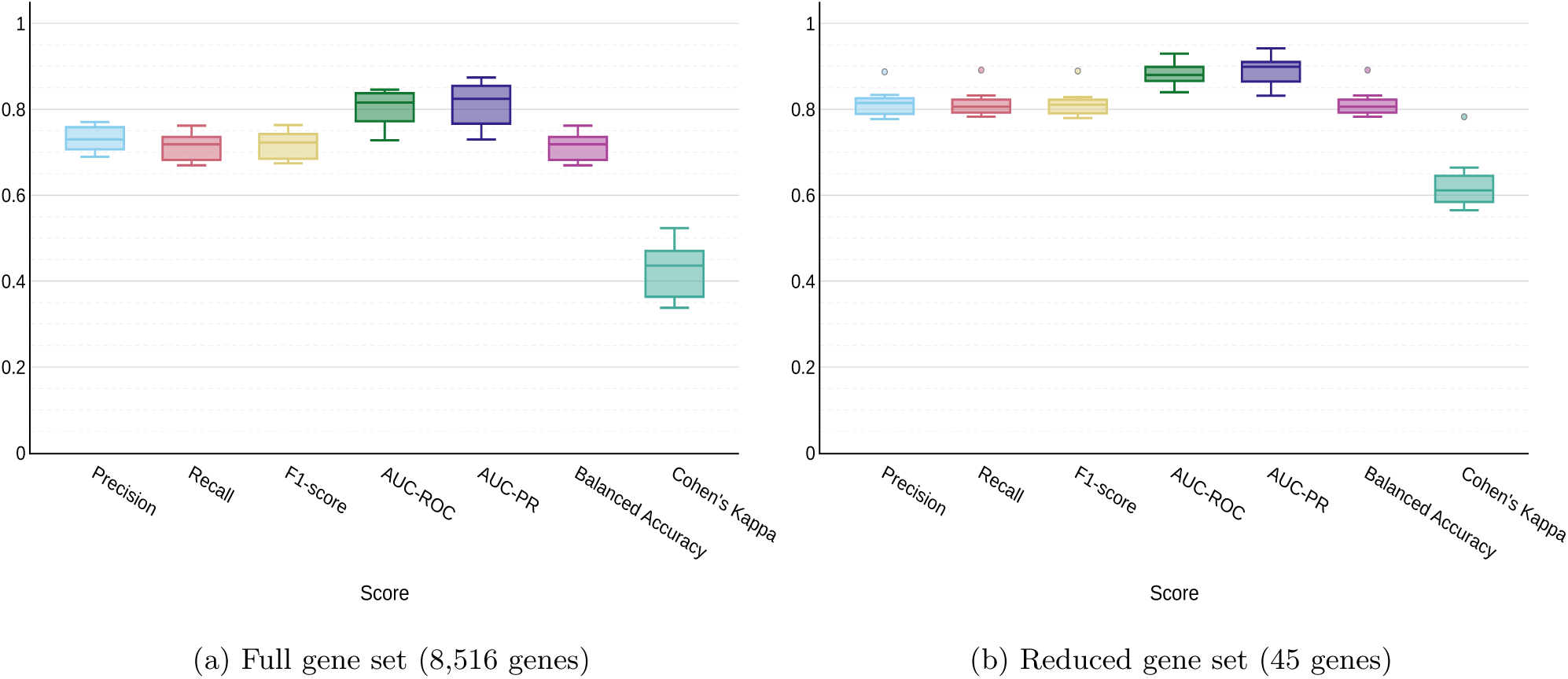
Early/late classification results on real patients. Performance metrics of an XGBoost classifier on early and late stages of ccRCC, evaluated with 10-fold cross-validation. Hyperparameters were optimized using Optuna. **a)** Classification using all 8,516 preprocessed genes. **b)** Classification using the refined set of 45 discriminative genes selected through multi-stage feature selection pipeline.

To deal with the high dimensionality of the gene expression data, we implemented a multi-stage feature selection pipeline that systematically identified the most informative genes while ensuring robustness (Methods 4.2). This approach yielded a final panel of 45 highly discriminative genes (Table 1). Retraining the classifier on this reduced panel, using the same training data and held-out test set, increased the mean AUC-ROC from the baseline 0.7877 to 0.8813 and improved the F1-score from 0.7156 to 0.8120 (Figure 2b).

**Table 1:**
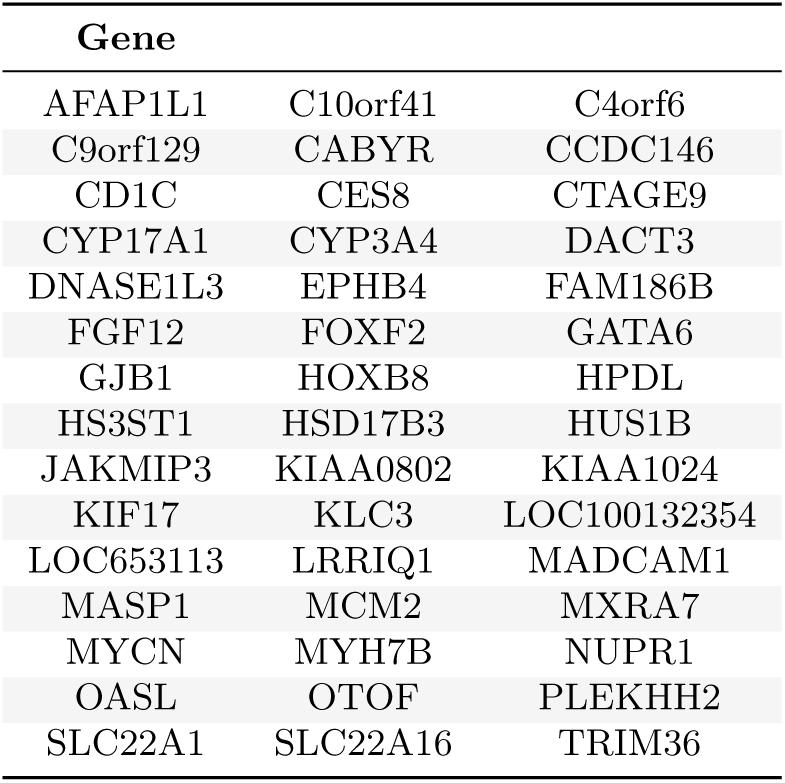
Top 45 discriminative genes for ccRCC stage classification. Genes identified through multi-stage feature selection pipeline, sorted alphabetically.

| Gene |  |  |
| --- | --- | --- |
| AFAP1L1 | C10orf41 | C4orf6 |
| C9orf129 | CABYR | CCDC146 |
| CD1C | CES8 | CTAGE9 |
| CYP17A1 | CYP3A4 | DACT3 |
| DNASE1L3 | EPHB4 | FAM186B |
| FGF12 | FOXF2 | GATA6 |
| GJB1 | HOXB8 | HPDL |
| HS3ST1 | HSD17B3 | HUS1B |
| JAKMIP3 | KIAA0802 | KIAA1024 |
| KIF17 | KLC3 | LOC100132354 |
| LOC653113 | LRRIQ1 | MADCAM1 |
| MASP1 | MCM2 | MXRA7 |
| MYCN | MYH7B | NUPR1 |
| OASL | OTOF | PLEKHH2 |
| SLC22A1 | SLC22A16 | TRIM36 |

To identify the biological processes associated with these 45 discriminative genes, we performed functional enrichment analysis using gProfiler (Methods 4.9). This revealed significant enrichment for pathways related to steroid and hormone metabolism, carnitine transport, and polyamine transport (Supplementary Figure S4).

### 2.2 Trajectory inference with VAE

The preprocessed ccRCC bulk RNA-Seq data (Methods 4.1) was split 80/20%, stratified by cancer stage, into training and test sets for a Variational Autoencoder (VAE; Methods 4.4). The VAE learns a latent representation of patient gene expression profiles through an encoder-decoder architecture: the encoder maps input profiles into a latent space, and the decoder reconstructs expression profiles from points in that space. This allows the decoder to generate synthetic gene expression profiles at intermediate points between two patients’ embeddings, which is the basis for the trajectory inference described below. The train/test split was preserved throughout all downstream analyses to prevent data leakage, ensuring no test-set information influenced representation learning, model development, or evaluation. We performed an architecture search to determine the best-performing VAE, in terms of minimizing the reconstruction error. To prevent the KL term from vanishing (posterior collapse) without imposing a strict target value, we used a beta-cycle-annealing schedule (Methods 4.4). The optimal VAE contained two layers of neurons at the encoder and decoder, generating a latent space of 256 dimensions (Supplementary Figure S6 and Supplementary Table S4). VAE models with more layers did not lead to better performance (Supplementary Results 4 and Supplementary Figure S7). The trained VAE showed a residual reconstruction error on the ccRCC dataset (Supplementary Figure S6, Supplementary Table S4). To correct this, we introduced a second-stage feedforward neural network trained directly on the VAE’s decoded output (Methods 4.5), following a similar data-space refinement strategy as [35]. This substantially reduced the reconstruction error (Supplementary Figure S8), aligning synthetic samples more closely with the empirical data manifold. Using the *sdmetrics* Python package, we found close statistical agreement between real and reconstructed test-set feature distributions across three metrics (Boundary Adherence, KS Complement, Correlation Similarity; Methods 4.5, Supplementary Table S5). The trained models reconstructed well almost all genes, except for 109 out of 8,516 (1.28%), which are lower-expressed (median 2.16 vs. 6.72 log_2_ units) and more often undetected (undetected in 5.8 times as many patients, on average) than the retained genes (two-sided Mann–Whitney U, *p* = 3.7 × 10*^−^*^40^ and *p* = 4.0 × 10*^−^*^52^, respectively; Supplementary Figure S9). This process allowed us to generate synthetic samples that closely resembled real patients.

To construct source-target pairs for trajectory inference, early- and late-stage patients were matched within sex and TCGA-reported ethnicity to minimize demographic confounding. Each early-stage patient was paired with multiple late-stage patients within their stratum, yielding a family of plausible trajectories rather than a single deterministic progression path (see Methods 4.6). To infer each trajectory, we linearly interpolated in latent space between the encoded source and target samples, and then decoded the intermediate latent points to obtain gene-expression profiles along the progression (details in Methods 4.6; see Discussion for alternative interpolation approaches). We chose to refer to VAE-inferred points as pseudo-time, given they do not correspond to real time points but reflect a time-dependent progression between stages. This resulted in a total of 269 source-target patient pairs between test-set patients, and 1,225 between train-set patients, each pair tracing the evolution of all 8,516 genes for 50 pseudo-time points (the first one being the source patient and the last one the target). Given the immense volume of data we chose to show an example trajectory between two patients in Figure 3a. The points at the leftmost and rightmost sides of the plot correspond to the early- (source) and late-stage (target) patients, respectively. The lines correspond to the VAE-inferred gene expression levels, showing rather complex patterns along the trajectory. As a set of control trajectories, we linked each patient at an early stage to itself, adding noise at the intermediate points (Supplementary Figure S11). These control trajectories served as the negative case of no transition between stages, remaining within an early-stage tumoral state. This design addresses a clinically relevant question directly: once a lesion has been diagnosed, can its subsequent evolution be distinguished from an indolent course? This reflects a central decision point in active surveillance for early-stage ccRCC, where serial imaging monitors localized tumors and intervention is deferred until clinically meaningful progression emerges, the key challenge being to distinguish indolent lesions from those showing growth kinetics or stage progression that would justify treatment [36, 37].

**Fig. 3:**
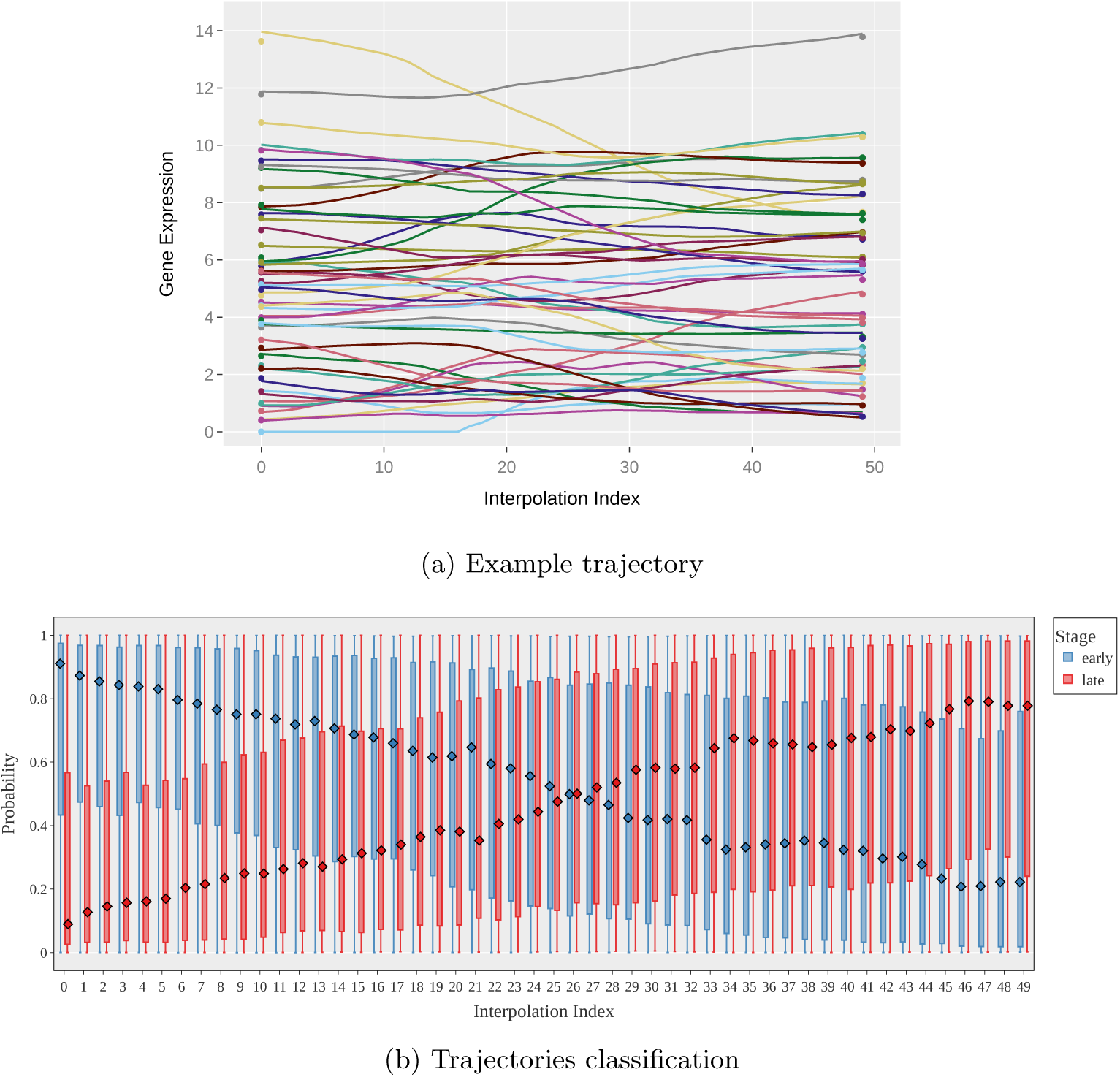
Synthetic trajectories between early- and late-stage ccRCC patients. **a)** Example trajectory between an early- and a late-stage patient showing gene expression levels of the 50 genes most important for stage classification. The leftmost and rightmost points correspond to the real early- (source) and late-stage (target) patients, respectively. Intermediate points were generated by linear interpolation in the latent space of the VAE, then reconstructing gene expression profiles. **b)** Classification of early and late stages along all synthetic trajectories. Distribution of probabilities assigned by the best-performing XGBoost classifier to synthetic patients at each point of the interpolated trajectories, showing the gradual transition from early-stage (left, high probability) to late-stage (right, high probability) classification.

Because TCGA-KIRC lacks ground-truth longitudinal trajectories, we separately validated our VAE-based trajectory-inference approach on a mouse CNS development dataset with genuine longitudinal structure, consisting of four real developmental time points per embryo [15] (Supplementary Methods 4). Using a leave-one-context-out scheme, we withheld each full developmental trajectory in turn, inferred it via latent-space interpolation between the remaining time points (as for the ccRCC case), and compared the result to the true, withheld time points. Across all four developmental stages, VAE-inferred profiles were substantially closer to the real data than a linear interpolation baseline in the original expression space (mean MSE 0.32–0.91 vs. 0.55–2.77; Cohen’s *d* = 0.81–2.63; Supplementary Result 4, Supplementary Figure S5), supporting the validity of applying this same trajectory-inference strategy to the ccRCC cohort, where no such ground truth is available.

To assess the quality of our synthetic trajectories, we applied the XGBoost classifier, mentioned above, trained solely on real ccRCC samples, to each point of the synthetic trajectories. At each VAE-inferred pseudo-timepoint, XGBoost estimated the probability of the given expression profile for all early-to-late stage trajectories to be either early or late stage, returning a per-stage distribution of probabilities. Figure 3b shows how the first points in the trajectories are mostly classified as early stage. The early stage probability gradually decreases along the trajectory, while the probability of late-stage classification increases. At the middle of the trajectories, there is a crossover point where the classifier assigns similar probabilities to both stages, and finally the majority of the patients are determined to be late-stage. As shown above, the test-set trajectories (Figure 3b) provide an independent validation of the classifier’s response to synthetic trajectories. As expected, the classification shift was even sharper in the trajectories of the train-set patients (Supplementary Figure S12). The control trajectories, used as a negative case with no stage transition, were classified almost uniformly as early stage (Supplementary Figure S13). Finally, the same classifier was trained on randomly shuffled labels to reflect a naive base-rate baseline classified nearly all points as early stage, i.e. the majority class, regardless of trajectory position (Supplementary Figure S14). Together, these results support that the stage transition captured by our trajectories reflects a genuine signal rather than an artifact of the classification procedure itself. The gradual classification probability shift along the trajectories from early to late stage indicates that the synthetic data may reflect meaningful stage transitions.

### 2.3 Stage Transition Forecasting

The first step to disease progression forecasting was to learn an interpretable, symbolic stage transition model in the form of an SFA, capturing gene expression-level patterns that govern stage transition in the underlying data. For that we used the ASAL (Answer Set Automata Learning) framework [33], an SFA induction system based on Answer Set Programming (ASP) [38]. ASAL learns SFA from possibly multivariate, positive and negative traces (i.e., event sequences), which, in our case, correspond to the positive (early-to-late stage) and the negative (control) multivariate gene trajectories generated by the VAE, for the top gene panel that was identified as explained above (Results 2.1).

ASAL encodes an SFA as an answer set program (i.e. a logic program in the ASP semantics) that combines a generic SFA interpreter, i.e., a sub-program that declaratively specifies how an SFA processes input sequences, a specification of the automaton’s structure (states and transitions), base predicates defined over the event tuples in the input sequences, and transition guard rules expressed as Boolean combinations of background predicates. This formulation makes both the automaton structure and the guard definitions learnable through the strong connections of ASP to symbolic learning. In particular, ASAL casts SFA induction as an abduc-tive, constraint-driven learning process: temporal structure and guard definitions are generated and tested against constraints related to predictive accuracy and model compression/minimality, in an effort to learn an optimal model that accepts the positive and rejects the negative training sequences, while being as simple/concise as possible, where simplicity is measured be the total number of states and transitions in the learned SFA and the number of predicates in the transition guard-defining rules. To learn a model with ASAL we further reduced the input dimensionality (i.e., the number of genes used) via SHAP-derived feature importance in an XGBoost classifier for discriminating between the source and end point in the trajectories. We used the ten most important genes identified by this classifier, with their expression levels discretized in a number of symbols (bins) to learn an SFA with ASAL, aiming to learn stage progression models with transition-enabling conditions expressed as Boolean combinations of gene expression thresholds for the different genes.

A learned SFA, achieving a test *f*_1_-score of 0.98, is illustrated in Figure 4. Because ASAL encodes transition guards as explicit, human-readable rules rather than opaque model weights, the learned SFA are, in principle, both auditable by clinical experts against known ccRCC biology and directly editable using domain knowledge.

**Fig. 4:**
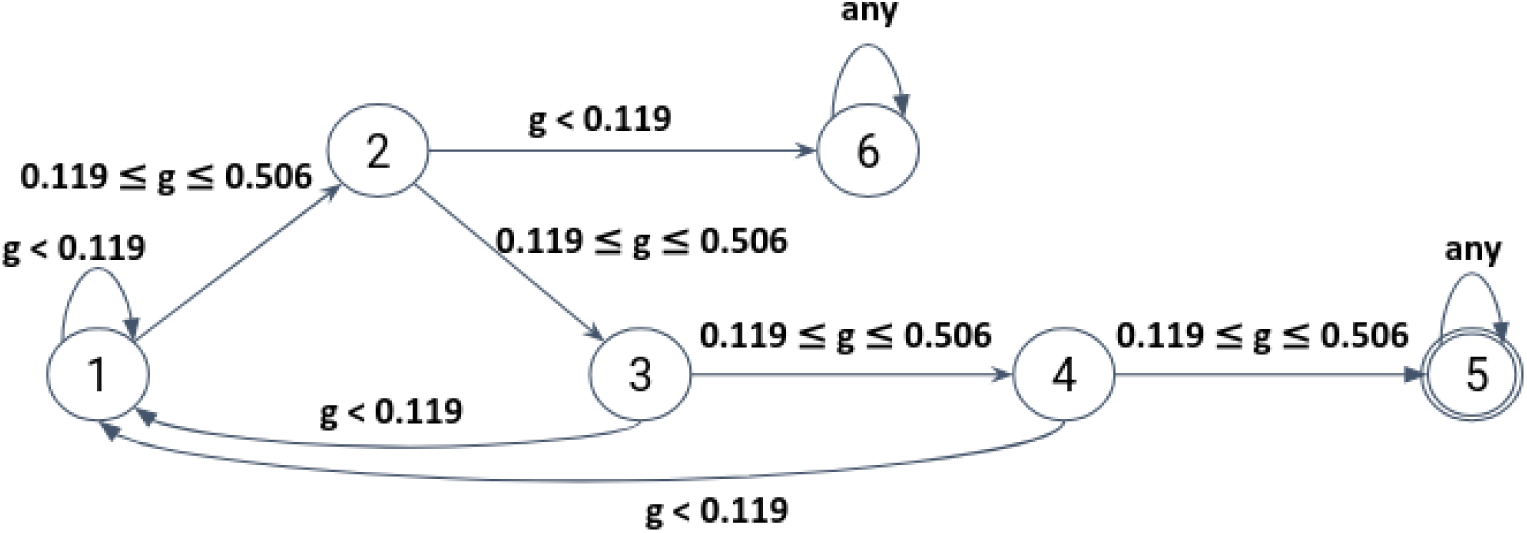
Learned Symbolic Finite Automaton (SFA) for ccRCC stage transition. An example SFA learned by ASAL from the panel of most informative genes identified by SHAP. State 1 is the start state. State 5 is the accepting state and state 6 is an explicit reject state. Both the accepting and the reject states are absorbing, i.e., they have no outgoing transitions. The gene *g* referenced in the SFA guards corresponds to the gene *SLC22A1*. The SFA expresses a sequential expression-level threshold pattern over this gene and represents a transition pattern dictating conditions that eventually lead to an early to late stage transition.

*SLC22A1*, ranked among the ten most discriminative genes for early/late-stage separation in our SHAP analysis (Methods 4.2), belongs to the SLC22 solute carrier family, which transports organic cations, anions, and drugs across cell membranes. Several other SLC22 family members show well-documented, stage-associated dysregulation in ccRCC [39–41]. Importantly, the behaviour identified by ASAL for *SLC22A1* indicates that the gene must rise to and sustain an intermediate expression level across several consecutive pseudo-time points for a trajectory to be recognized as a genuine stage transition. This example illustrates how our symbolic approach uncovers interpretable, temporally-resolved expression patterns rather than a single-timepoint threshold.

The learned patterns are input to Wayeb [34], a Complex Event Forecasting system. Wayeb is a probabilistic forecaster capable of predicting the potential occurrence of a declaratively defined complex event pattern within an event stream, before the event actually occurs, i.e. before the pattern is fully matched in the stream. In Wayeb, these patterns are typically defined as *Symbolic Regular Expressions*, or equivalently, SFA, such as those learned by ASAL. Wayeb works by estimating so called *waiting-time distributions*, i.e. probability distributions dictating the likelihood of reaching an accepting state in the input SFA from any other state in a number of steps. This allows Wayeb to emit probabilistic pattern completion forecasts and time-to-completion estimates while processing the input streams and moving across states in the monitored SFA. To model the statistical properties of the stream, Wayeb employs Variable-order Markov Models (VMMs), specifically, Prediction Suffix Trees (PST), which capture long-term dependencies, while avoiding the computational explosion associated with exhaustive enumeration in fixed-order models.

Figure 5 presents test-set forecasting performance, comparing the SFA-based Wayeb forecaster with a purely neural LSTM baseline for early classification. Wayeb reaches stable high performance by the 8th–10th point, while the LSTM reaches the *F* 1 *≥* 0.90 threshold earlier, at the 5th–6th point (10–12% of the sequence), reflecting its sensitivity to early discriminative signal. The LSTM also achieves a higher final macro F1-score (0.964 vs. 0.928), although the 3.6-point gap is modest.

**Fig. 5:**
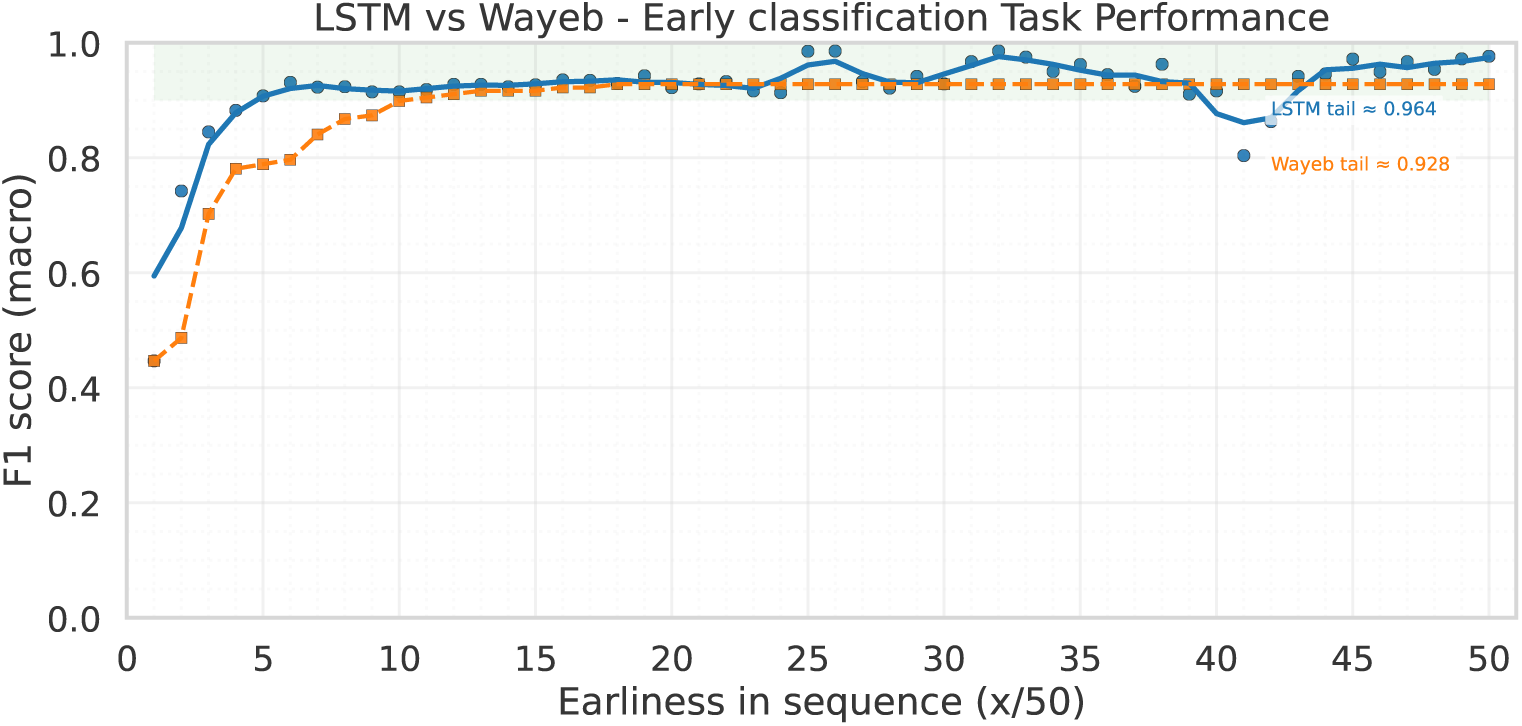
Stage transition forecasting results. Comparison of forecasting performance across synthetic trajectories (length 50 points) using Wayeb with the ASAL-learned SFA and a purely neural LSTM baseline for early classification. The symbolic approach (Wayeb) demonstrates comparable performance to the neural baseline, achieving a macro F1-score of 0.928 versus the LSTM’s 0.964, with a more gradual learning curve but providing full interpretability of the learned patterns and explicit confidence intervals for predicted transition timing.

This performance gap should be weighed against what the two approaches actually deliver. Unlike the LSTM’s single point score, Wayeb outputs a full probability distribution over transition timing at every step, together with an explicit, auditable SFA structure (Figure 4) rather than a distributed pattern encoded across network weights. As further addressed in the Discussion, these results indicate that Wayeb’s symbolic forecasts are close to LSTM-level accuracy while offering auditable predictions whose rules are structured for expert review, properties that matter directly for deployment in high-stakes clinical decision-making, where a system’s legibility can be as important as its raw predictive performance.

### 2.4 Biological Interpretation of Trajectories

We aimed to biologically interpret the synthetic trajectories by performing a pathway enrichment analysis along the generated pseudo-time in the test-set trajectories, comparing the forward trajectory timepoint with all the 72 solid tissue normal samples available from the TCGA-KIRC dataset. To do so, we performed a differential expression analysis (DESeq) followed by Gene Set Enrichment Analysis (GSEA) on all the gene sets available in Reactome (see Methods 4.10 for details). We considered only the significantly enriched pathways at each timepoint (FDR q-value *<* 0.05), displaying the sum of Normalized Enrichment Scores (NES) (see Methods 4.10) for all trajectories, for each pathway and timepoint. Figure 6 shows the top 50 pathways that change the most, in absolute terms, comparing the sum of NES values at the first and last timepoints. Supplementary Figures S16 and S17 show, on average, the top 50 most up-and down-regulated pathways, respectively. We observe that the different pathways tend to get upregulated with time more often than downregulated and a weak but significant positive correlation between the number of genes in each pathway and the absolute difference in NES between the first and last timepoints (Pearson’s *r* = 0.202 and *p* = 10*^−^*^25^). Kidney cancer is well-known as a metabolic disease[42, 43], reflected in Figure 6 by the differential enrichment of metabolic pathways (Biological Oxidations, Metabolism of Amino Acids and Derivatives, Selenocysteine Synthesis, Phase II - Conjugation of Compounds).

**Fig. 6:**
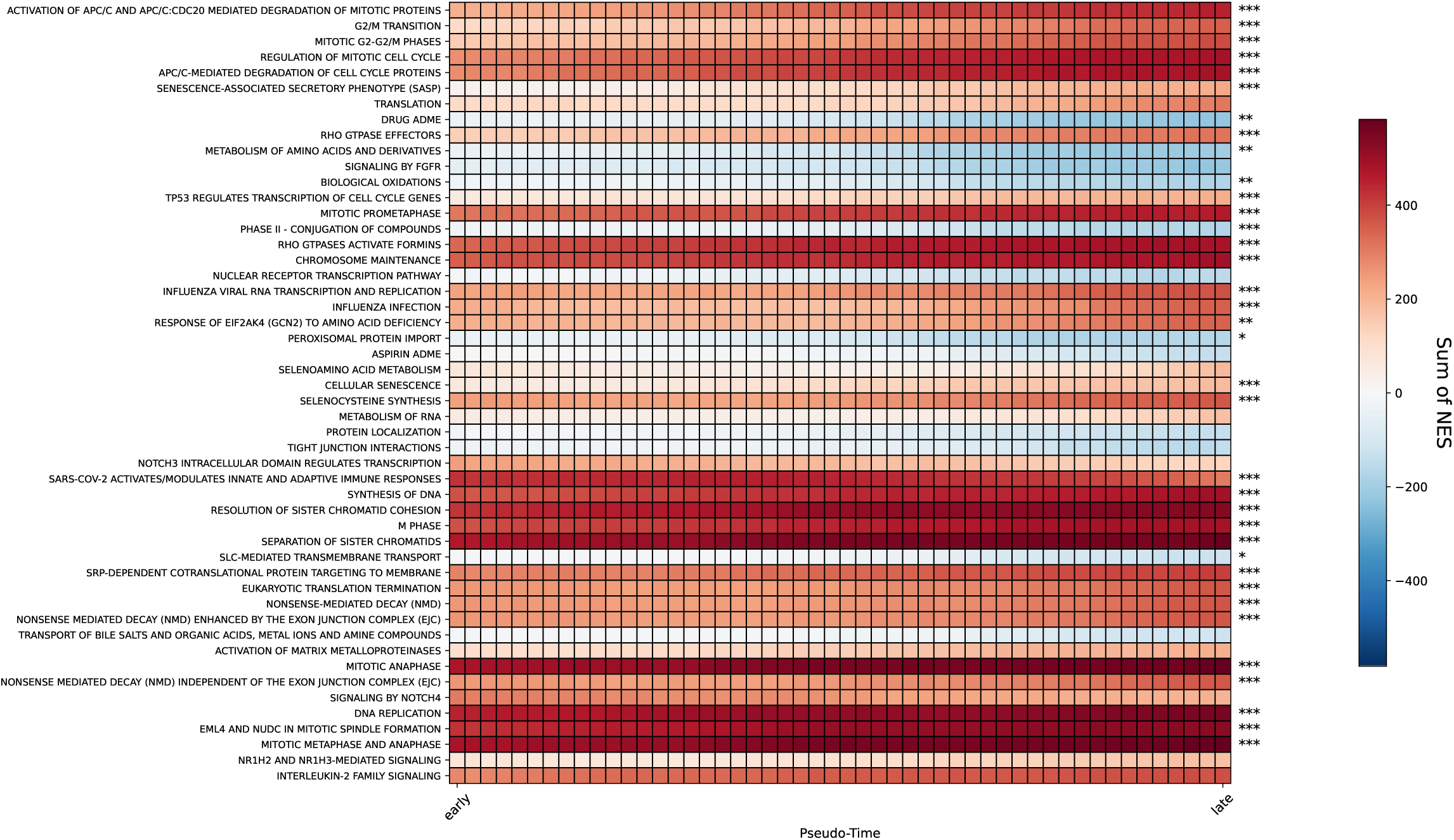
Gene Set Enrichment Analysis (GSEA) of synthetic trajectories. All pathways in Reactome were considered and a Differential Expression Analysis (DESeq) was run comparing test synthetic trajectories at each timepoint to normal tissue samples from TCGA, followed by a GSEA. Shown are the 50 pathways that change the most, in absolute terms, from the first to the last point of the trajectories. Red corresponds to upregulations, blue to downregulations, and white to no change in regulation. The equality of the distribution of the NES values between the first and last pseudotime points was tested using the Mann-Whitney U test. Significance levels are indicated as: *, *p ≤* 0.05; **, *p ≤* 0.01; and ***, *p ≤* 0.001.

Supplementary Figure S15a shows some relevant, literature-curated, pathways and their NES as the results of the GSEA on all the synthetic forward trajectories between test-set patients, while in Supplementary Figure S15b the NES has been min-max scaled between −1 and 1 to make relative changes across pseudotime more visually comparable across pathways. We find namely that Signaling by VEGF, related to angiogenesis, is largely upregulated throughout the disease progression. Some well-known ccRCC-related pathways [42–44] are also slightly altered, related to: the PI3K/AKT pathway (TP53 regulates metabolic genes), the Warburg effect (glycolysis), the TCA cycle, the SLITS and ROBOS pathway, apoptosis, DNA repair, and the metabolism (fatty acid metabolism). Besides, these pathways show a time-dependent behavior along pseudo-time. Apoptosis and the fatty acid metabolism pathways are among the few pathway showing a gradual downward trend with pseudotime. Meanwhile, DNA repair, regulation of expression of SLITS and ROBOS, and the citric acid (TCA) cycle and respiratory electron transport pathways become gradually upregulated. Some of them do show a gradual change in upregulation (DNA repair, Extracellular matrix organization, Metabolism of RNA) or downregulation (Drug ADME, Neuronal System, Protein Localization, Transport of Small Molecules) as patients get closer to the late stages and farther from the early stage. Our synthetic trajectories are then able to capture finer-grained changes among these important pathways with disease progression.

## 3 Discussion

In this work, we present a novel framework for modeling and forecasting longitudinal cancer stage progression from cross-sectional bulk gene expression data. The framework addresses two central limitations in the field: the scarcity of longitudinal cancer transcriptomic data, which we overcome through VAE-generated synthetic trajectories, and the lack of transparent, clinically auditable forecasting methods, which we address through Symbolic Finite-state Automata (SFA) and Complex Event Forecasting. By modeling ccRCC stage progression from these synthetic trajectories, and by forecasting impending transitions from partial observations, the framework moves beyond static case-control classification toward proactive, interpretable prediction of disease progression. To our knowledge, this represents the first application of this kind of hybrid AI to predict cancer stage progression.

Previous biomedical applications of VAEs have largely focused on learning low-dimensional representations of static transcriptomic data [10, 11], including the study of differences between early and late ccRCC stages [14]. Their use for synthetic data generation in biomedical omics remains comparatively underexplored, particularly for inferring temporal or longitudinal disease dynamics, a gap that we highlighted in a recent systematic review [16]. We address this gap directly, showing that a VAE trained on cross-sectional ccRCC profiles can infer plausible trajectories of disease worsening from early to late stage.

The generative abilities of VAEs in cancer biology remain underexplored, particularly in dynamic scenarios [16]; existing work on ccRCC stages has instead relied on static analyses, focusing on either classification [5] or differential expression [45]. By generating synthetic trajectories that mimic disease progression (Figure 3a), we circumvent this reliance on longitudinal data altogether, consistent with prior evidence that simple latent-space interpolation can generate complex, non-trivial transitions along a real data manifold [46].

We validated these trajectories in two complementary ways. First, an independent classifier trained only on real, static ccRCC patients, avoiding any data leakage (Supplementary Figure S18), correctly tracks a gradual shift in predicted stage as pseudo-time advances (Figure 2, Figure 3b), and the genes underlying this classifier are associated with steroid and hormone metabolism, carnitine transport, and polyamine transport (Table 1, Supplementary Figure S4), consistent with prior stage-classification studies in ccRCC [5, 8]. Second, pathway enrichment analysis (Figure 6) recovers stage-dependent changes in pathways already known to be altered in ccRCC, particularly metabolic processes, which is unsurprising given that kidney cancer is a well-established metabolic disease [42, 43]. Together, these two independent lines of evidence support that our synthetic trajectories replicate biologically plausible disease progression.

Beyond validating the synthetic trajectories, a central contribution of this work is forecasting the stage transitions they encode, and here the choice of forecasting method carries its own trade-offs. The LSTM, a widely used approach for sequence forecasting, outputs a single point score rather than a distribution over outcomes. Moreover, deep neural networks in general are well documented to be prone to poor calibration and overconfidence, even as their raw accuracy improves [26]. Wayeb, by construction, outputs a full probability distribution over transition timing at every step, derived from explicit waiting-time estimates rather than a black-box activation. In a clinical forecasting setting, this distinction is not incidental, as models that convey how much a prediction should be trusted, not just what the prediction is, are recognized as important for safely deferring low-confidence cases to human judgment in medical decision support [25]. The observed modest drop in aggregate F1-score is a reasonable cost for gaining access to this richer, probabilistic output.

Complementing this, the SFA underlying Wayeb’s forecasts is fully inspectable. Every transition guard is an explicit, human-readable condition over gene-expression thresholds (Figure 4), rather than a distributed pattern encoded across LSTM weights. This allows domain experts to directly audit whether a discovered pattern is biologically plausible, to reject or revise rules that are not, and to inject prior knowledge into the model without retraining, none of which is possible with the LSTM baseline. This kind of built-in interpretability has been argued to be preferable to post hoc explanation of black-box models, particularly for high-stakes decisions where the fidelity of an explanation to the model’s actual reasoning cannot be guaranteed [24]. We view this as the primary practical advantage of the symbolic approach: not that it achieves superior aggregate accuracy in this proof-of-concept setting, but that its errors and confidence are more interpretable and amenable to correction than those of a neural forecaster.

Our approach has several limitations. Because our external-classifier validation strategy relies on stage being well-separated in the real, static data, its diagnostic power is itself contingent on this separability. Among the ten cancer types, ccRCC showed the clearest early/late separation (Supplementary Figure S2). For cancers with weaker separability, a flat or noisy classifier response along a synthetic trajectory would be uninterpretable, it could reflect a genuinely poor trajectory, or simply a classifier with little signal to detect one. Extending this framework to other cancers therefore requires either stronger static classifiers or an independent validation strategy not bottlenecked by the same separability assumption. Similarly, our biological interpretation of the trajectories is currently limited to GSEA, while experimental validation, best pursued by dedicated wet-lab groups, remains an important direction for future work. A separate set of limitations concerns the forecasting stage specifically. The discretized, rule-based abstractions underlying the SFA may discard fine-grained temporal and quantitative information, likely contributing to its slightly weaker early predictive performance relative to the neural LSTM baseline. Symbolic abstractions are also often expected to generalize more robustly than purely neural models under distribution shift, since neural models are prone to latching onto spurious, dataset-specific “shortcuts” that fail to transfer once the input distribution changes, even when those shortcuts achieve strong in-distribution performance [47]; however, we could not test this directly, since training and test trajectories in our setting are both generated from the same underlying TCGA-KIRC distribution, preprocessing, and VAE-based construction procedure.

Future work can extend this framework along several directions. Because cancer progression cannot be fully captured by transcriptomics alone, integrating multi-omics data may reveal finer mechanistic detail, and incorporating single-cell data could help resolve the heterogeneity of the tumor microenvironment [48], building on a recent uptick of interest in generative, optimal-transport-based modeling of single-cell dynamics [49, 50], provided the focus remains patient-centric to preserve clinical relevance. The generative model itself could also be improved, for instance through normalizing flows [51] or training objectives that explicitly account for time-dependencies [52]. On the interpolation strategy specifically, we used linear interpolation in latent space, favoring an interpretable, minimal-assumption approach. Spherical linear interpolation (SLERP) or more sophisticated approaches, such as geodesic interpolation, which better respects the latent space’s probabilistic geometry, or optimal-transport-based trajectory inference [49, 50], could further improve the fidelity of intermediate synthetic profiles. On the forecasting side, end-to-end neurosymbolic architectures, or hybrid continuous-symbolic models with soft predicates or differentiable automata, may help better balance interpretability with early predictive accuracy.

We believe this work lays the groundwork for future research on cancer stage progression by combining generative and symbolic components to produce trustworthy synthetic data and clinically actionable forecasts as more detailed longitudinal datasets become available.

## 4 Methods

### 4.1 Data Collection and Preprocessing

The dataset used in this study contains the bulk RNA-Seq gene expression data from the TCGA database, downloaded through the [UCSC Xena browser](https://xenabrowser.net/). Genes were filtered out according to two criteria: genes with zero expression in at least 20% of the patients and genes whose expression mean and variance were below 0.5 *log*_2_ (*RSEM* + 1).

The same portal also provides clinical data, containing the cancer type, sex, and race information, as well as the cancer stage, for each patient. We found the top 10 most common cancer histological types in the TCGA database (Table S1) that included stages I, II, III, and IV patients (Supplementary Table S2). The stages were grouped as follows: I and II into “early” and III and IV into “late” stages. We trained an XGBoost classifier for 100 boosting rounds to distinguish between these two stages, using the gene expressions as features, and found the best performance was achieved in the case of ccRCC, also known as KIRC (Supplementary Figure S2). Before training the VAE, the ccRCC gene expression data were further preprocessed through outlier genes identification and their consequent removal using the Mahalanobis distance [53] as criteria. After preprocessing, the final dataset used for training and testing the VAE and XGboost models specific for ccRCC contains expression values of 8,516 genes in 530 patients.

### 4.2 Stage classification using static data and feature reduction

We employed the gradient boosting framework XGBoost (XGBClassifier) to develop a machine learning classifier that distinguishes between early and late-stage ccRCC. The dataset was partitioned into training (80%) and testing (20%) sets using stratified sampling to maintain class proportions. Within the training set, we implemented 5-fold cross-validation to optimize hyperparameters and assess model performance stability. To address the moderate class imbalance, we utilized balanced class weights via a parameter in XGBoost, that allows for calculating a balance between positive and negative weights. We optimized several key hyperparameters of the model using the Optuna framework with 100 trials, targeting maximization of F1-score.

To address the high dimensionality of the gene-expression data we implemented a multi-stage, model-guided selection pipeline. We preserved the original train/test partition of the VAE and split the training set further (stratified) to create an internal validation set. All selection decisions were made using only training/validation data so the held-out test set remained untouched. First, an XGBoost classifier was fit on the training partition to provide a baseline model. To obtain a robust, generalizable feature ranking we computed SHAP values with a TreeExplainer and aggregated importance as the mean absolute SHAP per feature averaged across training folds (stratified k-folds). Features were ranked by this averaged importance and we evaluated candidate top-k subsets by training a new XGBoost on the training partition and measuring macro-F1 on the validation partition. The 45 genes with the best validation macro-F1 were chosen. The final model was then retrained on the combined train and validation set using only the selected genes and evaluated on the untouched test set. We report macro-F1, accuracy, precision and recall on that held-out data.

### 4.3 Gene Expression Analyses

To determine Differentially Expressed Genes (DEGs) between the early and late stages, two consecutive statistical tests were performed. A Mann-Whitney U test was used to determine if the expression of genes was significantly different between the early and late stages. A p-value threshold of 0.01 was used.

Given that the p-value depends on the sample size, the effect size (Cohen’s d) was also calculated on the genes identified as differentially expressed. The *effsize* R package was used to calculate the effect size and its 99% confidence interval. We followed general guidelines to determine the importance of the effect size [54] as small (0.2 *< d ≤* 0.5), medium (0.5 *< d ≤* 0.8), and large (*d >* 0.8).

An analogous analysis to determine DEGs was performed, using the *limma* R package and empirical Bayes, and selected the DEGs as those with a p-value lower than 0.01. Both methodologies were compared to obtain the Supplementary Dataset 1, which contains the agreement on DEGs between both methods.

### 4.4 Training the Variational Autoencoder

We designed an unsupervised VAE [9] with two hidden layers in both the encoder and decoder. The encoder takes in as input dimension the number of genes, processes the data through a first hidden layer, and a second hidden layer estimates the mean *µ* and standard deviation *σ* of each dimension in the latent space. The reparametrization trick [9] was used to sample from one normal distribution *N* (*µ, σ*) per latent space dimension. The decoder then takes the sampled latent space, processes it in a complementary manner to that of the encoder, producing outputs in the original bulk-RNASeq gene expression data space.

The preprocessed data was split into training and testing sets using an 80/20% ratio, stratified by patient stage, and normalized using the MinMaxScaler function from the scikit-learn library. The VAE was trained with the Adam optimizer, using a learning rate of 10*^−^*^4^, a batch size of 8, and a linear *β*-cycle-annealing schedule [55] for the KL divergence loss. Consequently, the formula for the Evidence Lower Bound (ELBO) takes the form

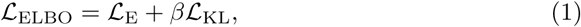

where *L*_E_ is the reconstruction loss, calculated as the Mean Squared Error (MSE), and *L*_KL_ is the non-negative KL divergence loss. The KL divergence loss was calculated between the approximate posterior and prior distributions,

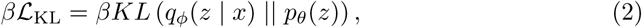

where *q_ϕ_*(*z | x*) is the approximate posterior distribution of the latent space *z* conditioned to the real space *x* given by a real but unknown set of parameters *ϕ*, *p_θ_*(*z*) is the prior distribution, and *θ* is the set of generative parameters which we want to estimate, i.e., the mean and standard deviation of each latent space dimension.

The schedule consists of three cycles of 200 epochs each. Within each cycle, the *β* parameter increases linearly from 0 to 1 for the first 100 epochs and remains constant for the next 100.

### 4.5 Synthetic Data Generation Pipeline

The trained VAE produced samples with a residual reconstruction error higher than desired for downstream use: the most optimal architecture showed a reconstruction of 220.51 as the average of the last 20 epochs in the test loss (Supplementary Figure S6, and Supplementary Table S4). In order to correct this, a post-processing, unsupervised neural network with four hidden layers was trained on the VAE-decoded data to minimize the reconstruction error between the real and decoded data. During training and testing, we kept the same patient train/test split as in the VAE model to avoid data leakage. The neural network was trained using the Adam optimizer with a learning rate of 10*^−^*^4^, a batch size of 8, for 1,000 epochs. The most optimal number of neurons for each hidden layer was obtained using *optuna* for 100 trials, and were set to 3,512; 824; 3,731; and 8,516 neurons (the last one coincides with the number of genes after preprocessing). This additional step yielded reconstructed data that closely resembled the real data, lowering the reconstruction error from 220.51 to 1.10 on average for the last 20 test epochs (Supplementary Figure S8). The post-processing step acts solely on the reconstruction error of the VAE, leaving the VAE’s KL-divergence intact.

To assess the quality of the synthetic generation, the real and a reconstructed test set data (effectively, this is synthetic data) were compared using the *sdmetrics* package in Python. We focused on three metrics to analyze the synthetic features (i.e., genes): Boundary Adherence, Kolmogorov-Smirnov (KS) Complement, and Correlation Similarity. The Boundary Adherence measures the frequency with which each feature in the synthetic data respects the minimum and maximum values of the real data feature distribution. The KS Complement computes the Kolmogorov-Smirnov (KS) statistic between each real and synthetic feature, and returns its complement (1-KS) so that 0 marks completely different distributions, and 1 marks identical distributions. The Correlation Similarity compares the real and synthetic Pearson correlation for each pair of features, then compares them to get a score between 0 (completely different correlations) and 1 (identical correlations).

### 4.6 Trajectory Inference with the VAE

Synthetic patients were generated at intermediate time points between patients at the early and late stages. To do so, a network was constructed using the early-stage patients as sources and the late-stage patients as targets, distinguishing between train-set and test-set patients. Patients were stratified by sex (male and female) and ethnicity (White, Black or African American, and Asian), following TCGA’s denominations. For the two underrepresented ethnicities (Black or African American and Asian), we linked every early-stage patient to all available late-stage patients. For the remaining majority ethnicity (White), 5 patients were selected at random at the late stage for every early-stage patient.

To prevent data leakage between the trajectories used for VAE/classifier training and those held out for evaluation, we ensured that no patient anchor (source or target) was shared between a train-set trajectory and a test-set trajectory. This was enforced by constructing a graph in which patients are nodes and an edge connects any two patients linked as a source-target pair; each connected component of this graph was then assigned entirely to either the training or the test set, ensuring that no patient appearing in a training trajectory could also appear as an anchor in a test trajectory (Supplementary Figure S18).

Once the patients had been linked, their gene expression was embedded in the latent space of the trained VAE. There, a linear interpolation of 48 points was performed. These trajectories included a total of 50 timepoints when accounting for the source and target patients. The 50 points were decoded to the original space, returning to the 8,516 genes obtained after preprocessing.

### 4.7 Control Trajectory Inference

Control trajectories were generated to use as a negative case for forecasting that emulates a case of no progression. Instead of progressing from an early to a late stage, the control trajectories remained in the early stage. To generate these control trajectories, a simpler approach was followed. Every early-stage patient was linked to itself with a straight line of 50 time points in the real space of gene expression. At each intermediate time point, a Standard Gaussian Noise (mean 0 and standard deviation 1) was added to each gene expression value.

### 4.8 Classification of time points in synthetic trajectories

The synthetic trajectories were assessed by classifying all time points. The XGBoost classifier trained on the real data, which had only seen patients from the original, real train set split (avoiding data leakage), was applied to every point in the synthetic trajectories. This yielded a probability for each trajectory time point to belong to either the late or early stage in ccRCC. Altogether, each time point showed a distribution of probabilities across all trajectories. In this way, a classifier that had only seen the real, static data was used to validate the synthetic, dynamic trajectories inferred with the VAE.

### 4.9 Static Enrichment analyses

To analyze selected genes from feature selection (see Methods 4.2), enrichment analysis was performed using the *gprofiler2* package in R. The analysis aimed to identify significantly enriched biological terms and pathways among genes important for ccRCC early and late stage classification, as determined from our feature selection pipeline (Results 2.1 and Methods 4.2). The enrichment was performed on annotated databases, namely Gene Ontology (GO), Reactome (REAC), Kyoto Encyclopedia of Genes and Genomes (KEGG), and WikiPathways (WP). The results of the enrichment analysis (Supplementary Dataset 2) include terms that are significant in humans, with a p-value below 0.05, corrected for with false discovery rate, considering annotated genes.

### 4.10 Dynamic Enrichment analyses

In order to analyze the biological validity of the inferred synthetic trajectories, we built a pipeline based on Differential gene expression analysis (DESeq) followed by a Gene Set Enrichment Analysis (GSEA). This pipeline was applied to all synthetic trajectories, including all time points and genes. Given the large volume of data analyzed, High-Performance Computing was required to run the pipeline.

The pipeline consisted of the following steps. First, using the PyDESeq2 Python package [56], DESeq was performed on all trajectories, comparing each time point to all the tumor-adjacent samples (the controls for DESeq) present in the TCGA-KIRC dataset, returning a rank file of differential gene expression for each trajectory and time point. Second, a GSEA was run on each rank file, using as reference gene sets all the pathways from Reactome. GSEA produced a report, including, namely, the Normalized Enrichment Score (NES) and an FDR-corrected q-value for each pathway, at each trajectory and time point pair. Third, we selected the significant cases (FDR q-value *<* 0.05) and kept the NES values. Finally, the NES scores were summed across all trajectories, for each time point and pathway, to get an overview of the pathway enrichment dynamics for all trajectories (as seen in Figure 6).

## Supporting information

Supplementary Dataset 1. Significantly differentially expressed genes

Supplementary Dataset 2. Static enrichment analysis results

Supplementary Dataset 3. Dynamic enrichment analysis results

## 5 Code Availability

The code used in this study is available on GitHub: gprolcastelo/renalprog. The repository includes a detailed documentation, with instructions to reproduce the experiments herein: https://gprolcastelo.github.io/renalprog/. The symbolic automata learning and forecasting components rely on ASAL (https://github.com/nkatzz/asal) and Wayeb (https://github.com/ElAlev/Wayeb).

## 6 Data Availability

The preprocessed TCGA-KIRC dataset, the VAE-generated synthetic longitudinal trajectories, and the GSEA results are available on Zenodo with DOI 10.5281/zen-odo.17987300. Besides, the trained models are also available on the Hugging Face repository gprolcastelo/evenflow_models.

