## Supplementary Information for "Interpretable Forecasting of Kidney Cancer Progression via Generative AI and Symbolic Reasoning"

### Supplementary Material

#### Contents

Supplementary Datasets . . . . . . . . . . S1
Supplementary Results . . . . . . . . . . S1
Supplementary Methods . . . . . . . . . . S2
Supplementary Tables . . . . . . . . . . S4
Supplementary Figures . . . . . . . . . . S7
Supplementary References . . . . . . . . . . S23

#### Supplementary Datasets

**Supplementary Dataset 1. Significantly differentially expressed genes.** Differentially expressed genes between early and late stages in the real dataset, with a p-value lower than 0.01. The genes included correspond to the agreement between Limma and Mann-Whitney U test (see Methods 4.3). The table includes the p-values for each method, as well as the log fold change (logFC) calculated with Limma, and the effect size (Cohen's $d$) for the Mann-Whitney U test.

**Supplementary Dataset 2. Static enrichment analysis results.** This dataset includes the terms and pathways that were found to be significantly enriched. The analysis was performed with gprofiler on the genes determined to be important for classification of the ccRCC stages (see Methods 4.9).

**Supplementary Dataset 3. Dynamic enrichment analysis results.** This dataset includes the results of the Differential Expression analysis followed by GSEA on the synthetic trajectories (see Methods 4.10). Columns correspond to the inferred time points, 0 being the real patients at the early stage, and 49 to the real patients at the late stage. Rows correspond to the 2,630 pathways available in the Reactome database.

#### Supplementary Results

##### Mouse CNS development dataset

The TCGA-KIRC dataset lacks clearly defined trajectories, as it only contains independent samples from patients at different stages. Hence, our first step was to validate our VAE-based Synthetic Data Generation pipeline (Methods 4.5) approach on a suitable dataset. We resourced to a mouse CNS development dataset from [1] (described in Supplementary Methods ) and proceeded to train a VAE on a leave-one-context-out (LOCO) basis (see Supplementary Methods ), leaving out one full developmental trajectory at each training fold. As a further difference with the main paper, the post-processing network was optimized with *ax* [2] instead of *optuna*, as a refinement on the pipeline, providing a more advanced hyperparameter optimization. Besides, in the post-processing network we considered neurons dropout to prevent overfitting to this smaller dataset [3]. The optimal VAE was determined to have a hidden layer of 512 and 128 neurons at the encoder and complementary at the decoder, with the latent space size of 64 dimensions. The post-processing network had 3 hidden layers of 128; 128; and a last layer that varies between 8,896 and 9,282 neurons, coinciding with the number of genes after preprocessing for each LOCO fold. Just like in the main text, the trajectories were inferred by interpolating straight lines in the latent space between encoded samples, then decoding the intermediate points. We compared the inference of each day in the left-out context to its real counter part and a baseline that consisted on the same straight-line inference on the real space. Supplementary Figure S5 shows the Mean Squared Error (MSE) distribution between the real time points and the synthetic, VAE-generated points, as well as with the straight-line baseline. The difference between the real and inferred samples remained low in our VAE-based scenario, clearly beating the baseline at all days: mean MSE 0.91 versus 1.33 at E11 ($p = 5.4 \times 10^{-6}$, Cohen's $d = 1.04$), 0.32 versus 0.55 at E13 ($p = 3.9 \times 10^{-3}$, $d = 0.81$), 0.42 versus 1.35 at E15 ($p = 2.7 \times 10^{-6}$, $d = 2.63$), and 0.41 versus 2.77 at E18 ($p = 2.3 \times 10^{-12}$, $d = 1.27$). All effect sizes reflect a very large effect.

#### Larger VAE models do not lead to better performance

More complex VAEs were trained on the TCGA-KIRC cohort, with an extra hidden layer on the decoder, with the intention to lower the reconstruction error. However, the performance was actually worse than in the simpler model described on the main text. Figure S7 shows the losses of the VAE with the more complex decoder. Our result is in accordance with [4], who showed that increasing the model's complexity may lead to lower KL divergence, and, consequently, to the dreaded posterior collapse.

### Supplementary Methods

#### Mouse CNS development dataset

Bulk RNA-seq data was obtained from [1], which included already a preprocessed, normalized version of the data. The dataset contains the information of 64 samples from two mice embryos, with two different conditions: wild type (WT) and Eed-c knockout (KO); at four different developmental time points: E11.5, E13.5, E15.5, and E18.5; and from four different tissues: forebrain (fb), midbrain (mb), hindbrain (hb), and spinal cord (sc). We followed the same preprocessing steps as described in our main text Methods section 4.1 to remove low-variance and outlier genes.

#### Synthetic Mouse CNS trajectories

Our VAE-based pipeline (see Methods 4.5) was trained on the mouse CNS development dataset and generated synthetic trajectories, following the Methods described in the main text (4.4 and 4.5). Besides, in order to avoid data leakage and maximize training data in such a small dataset, we ran a leave-one-context-out (LOCO) training schedule. This consists on leaving one entire developmental trajectory for both biological replicates across all four days (e.g., both replicates of the four days in WT fb1). Synthetic trajectories were generated in the left-out context via interpolation in the latent space from every combination of two days, leaving the others out as references and measuring the error levels. The generative strategy was repeated for a straight-line-on-real-data baseline comparison. In both cases, one interpolated time point represents one real day. This way, the performance of the VAE can be assessed when generating synthetic trajectories at inferred time points. To compare the error between the VAE-based generation strategy and the baseline, we ran a two-sided Mann-Whitney U-test per each day error distribution (VAE-based vs. baseline) and calculated the corresponding Cohen's d to report effect sizes.

### Supplementary Tables

| Cancer Type | Histological Type | Count |
|---|---|---|
| BRCA | Infiltrating Ductal Carcinoma | 767 |
| KIRC | Kidney Clear Cell Renal Carcinoma | 533 |
| LUAD | Lung Adenocarcinoma | 511 |
| LUSC | Lung Squamous Cell Carcinoma | 500 |
| HNSC | Head & Neck Squamous Cell Carcinoma | 443 |
| BLCA | Muscle invasive urothelial carcinoma (pT2 or above) | 407 |
| COAD | Colon Adenocarcinoma | 380 |
| THCA | Thyroid Papillary Carcinoma - Classical/usual | 358 |
| LIHC | Hepatocellular Carcinoma | 343 |
| BRCA | Infiltrating Lobular Carcinoma | 200 |

**Supplementary Table S1**: **Top ten most common cancer histological types in the TCGA dataset.** The columns show the TCGA cancer abbreviation, the cancer full name, and the number of patients found in the dataset (raw pull).

| Cancer Type | Stage | Count |
|---|---|---|
| BLCA | I | 0 |
| | II | 130 |
| | III | 141 |
| | IV | 136 |
| BRCA | I | 163 |
| | II | 564 |
| | III | 222 |
| | IV | 18 |
| COAD | I | 65 |
| | II | 153 |
| | III | 104 |
| | IV | 58 |
| HNSC | I | 26 |
| | II | 74 |
| | III | 78 |
| | IV | 265 |
| KIRC | I | 268 |
| | II | 57 |
| | III | 125 |
| | IV | 83 |
| LIHC | I | 171 |
| | II | 83 |
| | III | 85 |
| | IV | 4 |
| LUAD | I | 278 |
| | II | 123 |
| | III | 84 |
| | IV | 26 |
| LUSC | I | 245 |
| | II | 163 |
| | III | 85 |
| | IV | 7 |
| THCA | I | 212 |
| | II | 29 |
| | III | 74 |
| | IV | 43 |

**Supplementary Table S2**: **Top ten most common cancer histological types in the TCGA dataset.** The table includes the number of patients at the different stage, for each cancer type.

| Method | Significantly DEGs |
|---|---|
| Limma | 3,218 |
| Mann-Whitney U | 3,321 |
| Both Methods | 2,979 |

**Supplementary Table S3**: **Comparison of Differentially Expressed Genes (DEGs) between early and late ccRCC stages.** Number of significantly DEGs identified by Limma, Mann-Whitney U test, and genes identified by both methods. Genes with p-value $p < 0.01$ were considered significant.

| | 16 | 32 | 64 | 128 | 256 | 512 |
|---|---|---|---|---|---|---|
| 256 | 244.215800 | 229.593448 | 224.419173 | 221.486604 | 227.734442 | 231.680966 |
| 512 | 253.749909 | 229.316439 | 221.484050 | 221.654277 | 220.512593 | 221.620340 |
| 1024 | 265.369299 | 241.539643 | 230.140319 | 223.672304 | 222.586957 | 253.853000 |
| 2048 | 257.397065 | 240.704352 | 231.497607 | 227.839528 | 245.937153 | 318.631503 |
| 4096 | 258.441426 | 245.624789 | 231.042961 | 231.736211 | 272.163110 | 361.652005 |

**Supplementary Table S4**: VAE losses for different model architectures. Rows show the number of neurons at the first hidden layer of the encoder (and last layer of the decoder). Columns show the latent space size. Values represent the average reconstruction loss of the last 20 epochs of testing.

| | mean | std | min | 25% | 50% | 75% | max |
|---|---|---|---|---|---|---|---|
| Boundary Adherence | 0.993 | 0.0452 | 0 | 0.991 | 1 | 1 | 1 |
| KS Complement | 0.846 | 0.0982 | 0 | 0.83 | 0.858 | 0.887 | 0.953 |
| Correlation Similarity | 0.941 | 0.0444 | 0.588 | 0.916 | 0.95 | 0.977 | 1 |

**Supplementary Table S5**: **Assessing synthetic data with sdmetrics.** The table shows the summary statistics of three metrics from the *sdmetrics* Python package: Boundary Adherence, KS Complement, and Correlation Similarity. We compared the test set of real patients with the test set of reconstructed patients using our synthetic-data generation pipeline. See Supplementary Figure S10 for boxplots of the distributions of each metric.

### Supplementary Figures

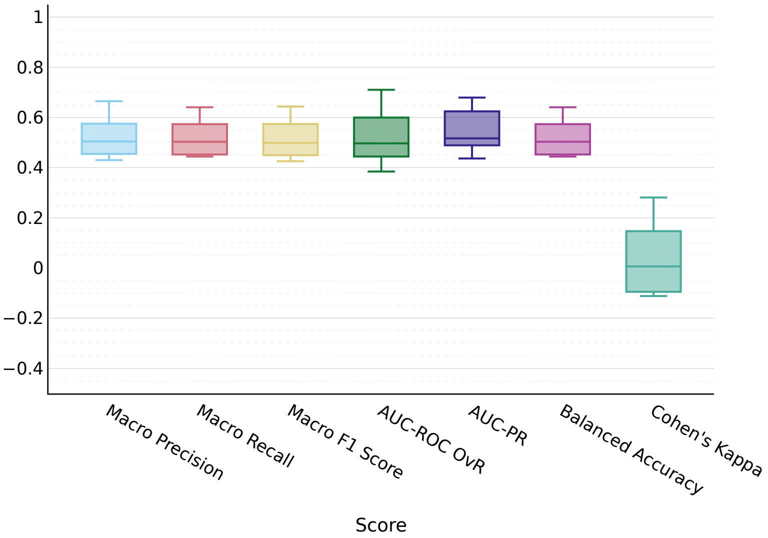


**Supplementary Figure S1**: **Randomization experiment on early and late stage classification of ccRCC patients.** The boxplot shows the performances of an XGBoost classifier on early and late stages of ccRCC, with 10 different seeds used to split the data into training and test sets. At each iteration, 'early' and 'late' labels were randomly allocated as patient stages, keeping the original number of samples per class. The classifier hyperparameters were optimized with Optuna. Results show this classification is random, as the median for all metrics is around 0.5, except for the Cohen's Kappa, around 0.

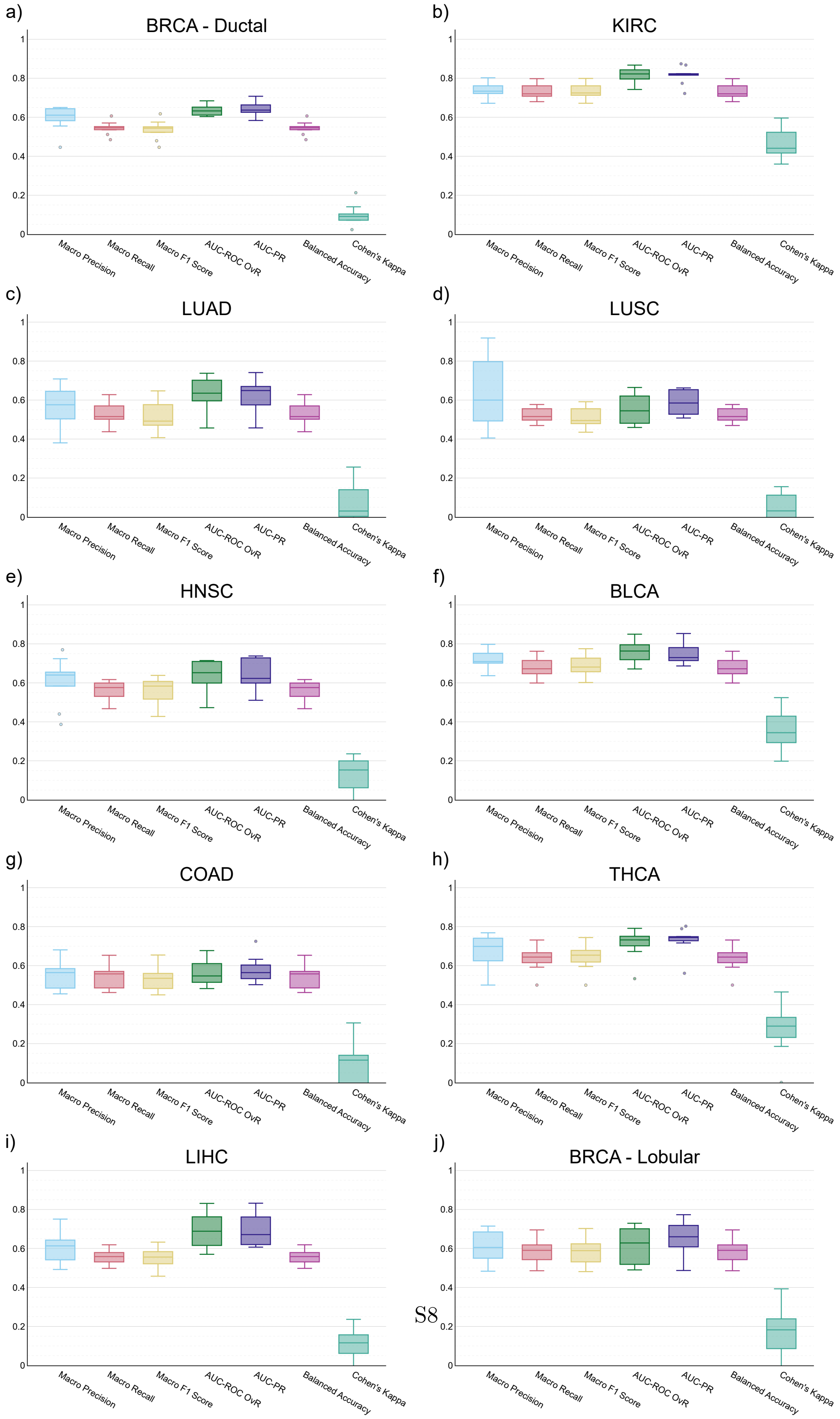


**Supplementary Figure S2**: **Classification on early and late cancer stages**. Top 10 most common cancer types in the TCGA dataset. **a)** Breast Invasive Carcinoma (BRCA) - Infiltrating Ductal Carcinoma, **b)** Kidney Clear Cell Renal Carcinoma (KIRC, also known as ccRCC), **c)** Lung Adenocarcinoma (LUAD), **d)** Lung Squamous Cell Carcinoma (LUSC), **e)** Head & Neck Squamous Cell Carcinoma (HNSC), **f)** Muscle invasive urothelial carcinoma (pT2 or above) (BLCA), **g)** Colon Adenocarcinoma (COAD), **h)** Thyroid Papillary Carcinoma - Classical/usual (THCA), **i)** Hepatocellular Carcinoma (LIHC), **j)** BRCA - Infiltrating Lobular Carcinoma.

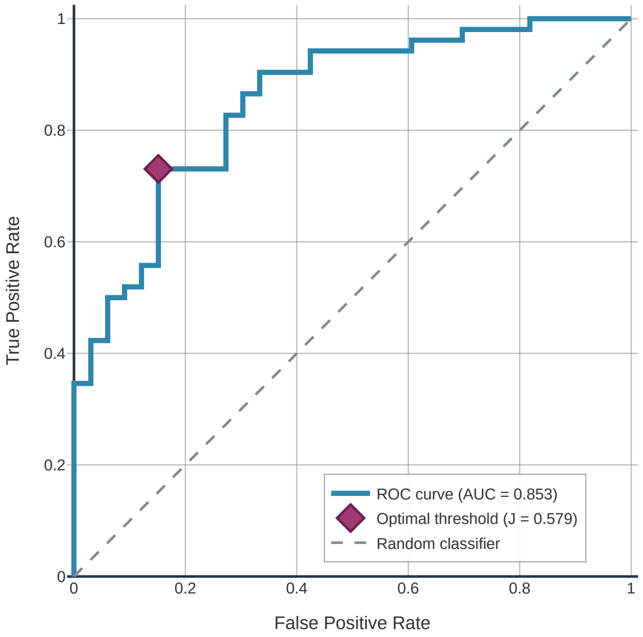


**Supplementary Figure S3**: **ROC curve of best-performing early/late ccRCC stage classifier, assigning early stage as the positive case.** The receiver operating characteristic (ROC) is shown for the best-performing XGBoost model to distinguish early and late stages in ccRCC. The x-axis represents the False Positive Rate (FPR) and the y-axis is the True Positive Rate (TPR). The area under the curve (AUC) is 0.853. Youden's J statistic is 0.579, the optimal threshold is 0.793, located at TPR 0.731 and FPR 0.152, is highlighted in the plot with a dark pink diamond.

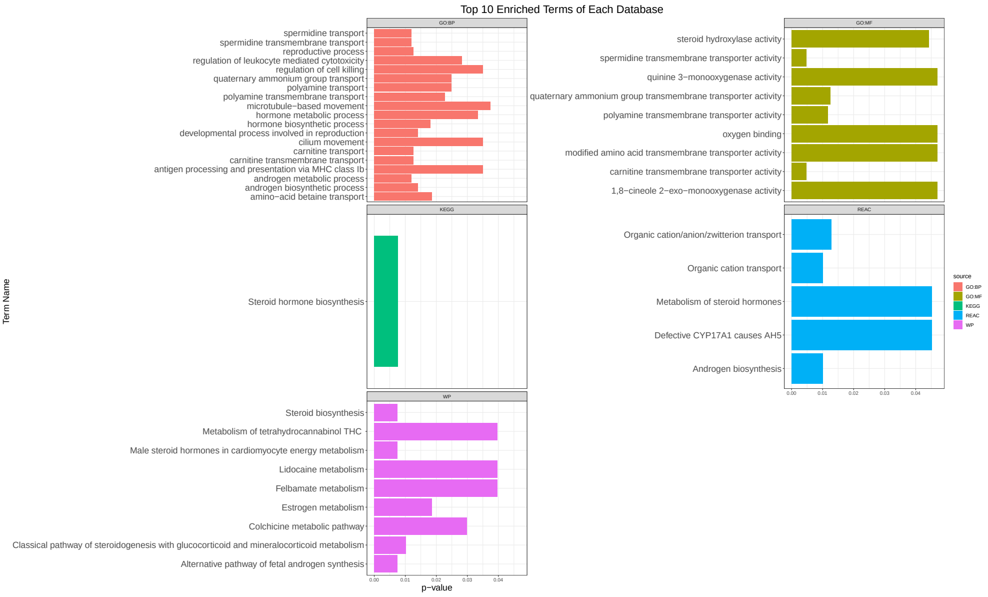


**Supplementary Figure S4**: **Enrichment analysis on genes that improve classification of ccRCC early and late stages**. The figure shows (at most) the top 10 most significant terms and pathways for each of the databases considered. Each plot is titled with the name of the corresponding database: Gene Ontology (GO), organized in two aspects, Biological Process (GO:BP) and Molecular Function (GO:MF); Reactome (REAC) (4); Kyoto Encyclopedia of Genes and Genomes (KEGG) (5); and WikiPathways (WP) (6). The enrichment analysis was conducted using gprofiler on the genes identified as important for the classification of ccRCC early and late stages.

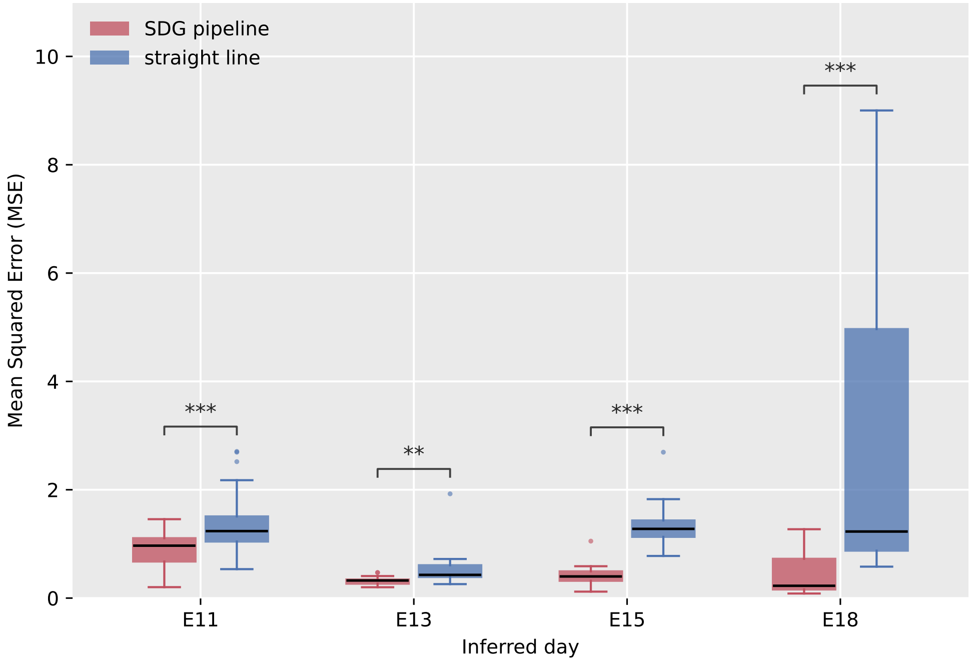


**Supplementary Figure S5**: **Mean Squared Error (MSE) between real and synthetic points in the mouse CNS dataset**. MSE corresponding at each inferred day were calculated between the real and the synthetic cases (our SDG, in red, and the straight line baseline, in blue), across all the leave-one-context-out (LOCO) folds. Each day distribution pairs were compared with a two-sided Mann–Whitney U tests: ** p ¡ 0.01, *** p ¡ 0.001. Cohen's d were 1.04, 0.81, 2.63 and 1.27, respectively.

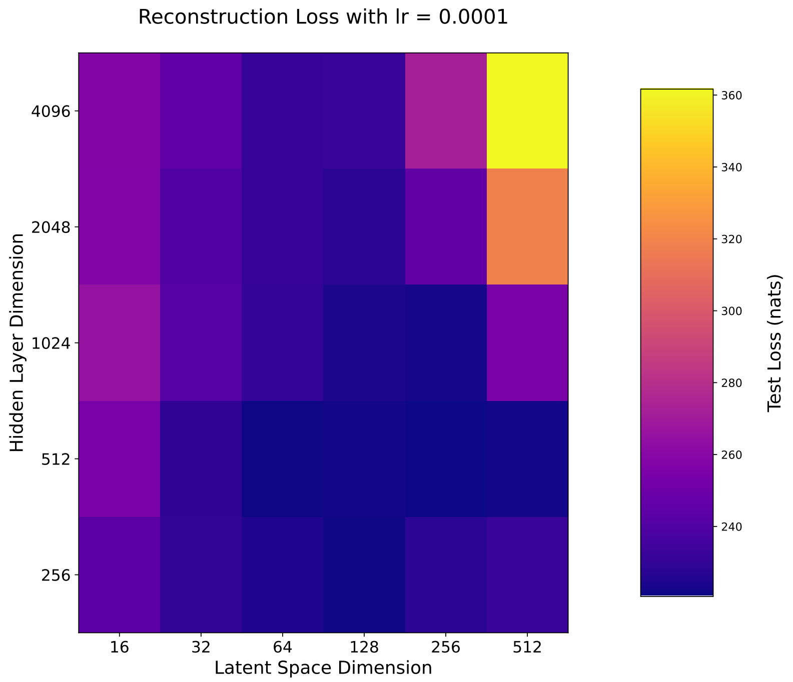


**Supplementary Figure S6**: **VAE losses for different model architectures.** Rows show the number of neurons at the first hidden layer of the encoder (and last layer of the decoder). Columns show the latent space size. Values represent the average reconstruction loss of the last 20 epochs of testing.

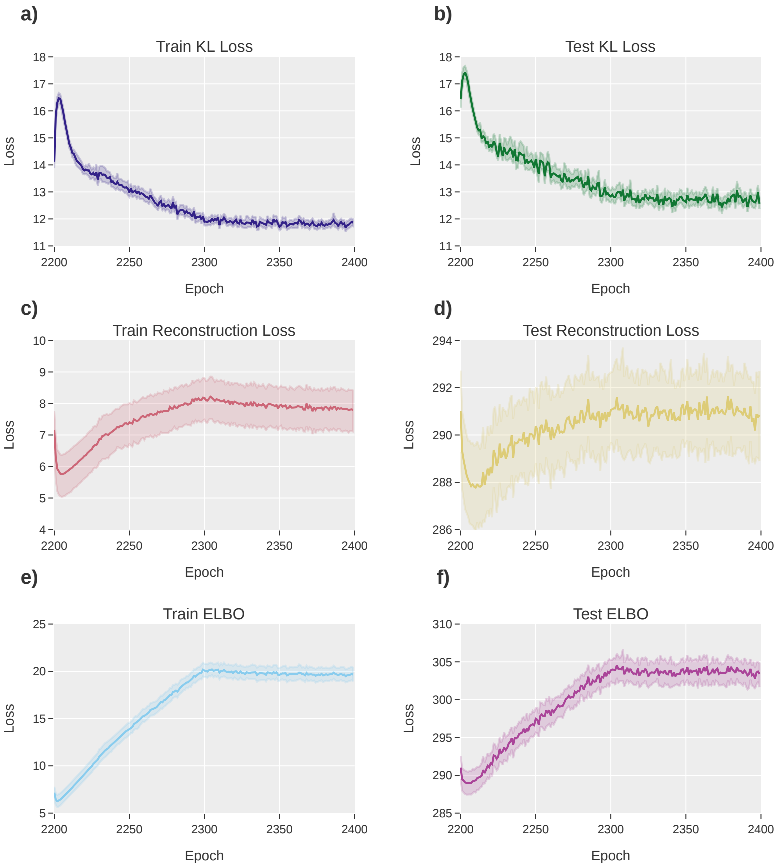


**Supplementary Figure S7**: **VAE testing losses when using more complex decoders**. **a,b)** KL divergence. **c,d)** Reconstruction Loss. **e,f)** Evidence Lower Bound (ELBO). The plots show the last of the annealing cycles for testing, which comprises the last 200 epochs of the process. We set an extra layer in the decoder, and ran 50 trials in optuna to reduce the reconstruction error. The solid lines represent the mean value of the trials and shaded areas represent the 99.9% confidence interval (CI). The reconstruction error is a lot larger when training a more complex decoder compared to that of the main paper. Besides, the model shows overfitting problems given the much larger reconstruction error (and, consequently, ELBO) in testing compared to training.

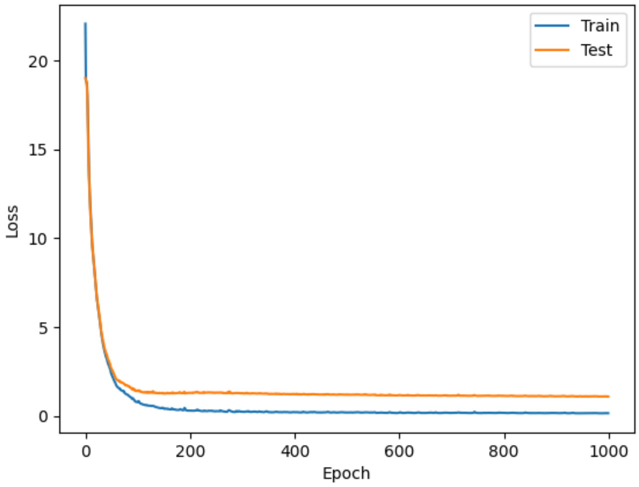


**Supplementary Figure S8**: Training (blue) and test (orange) losses for the post-processing step.

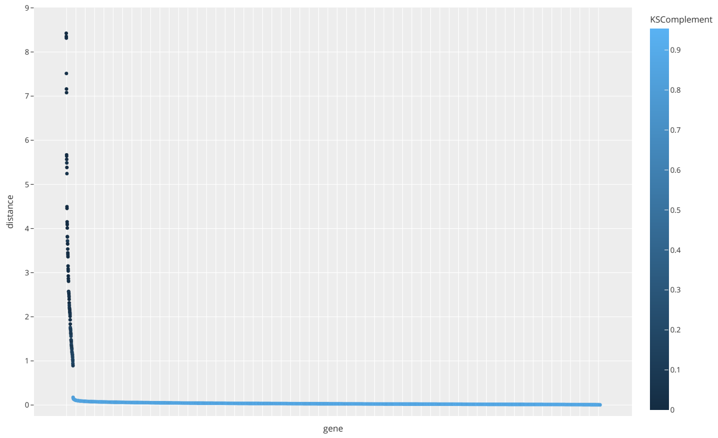


**Supplementary Figure S9**: **Gene expression reconstruction quality.** The y-axis shows the Wasserstein distance between the real and reconstructed gene expression distributions for each gene. The x-axis represents each individual gene, sorted by decreasing distance. The color of each point represents the complementary Kolmogorov-Smirnov (KS) statistic (1 - KS) or KSComplement between the real and reconstructed data. The most extreme values of KSComplement are 1 (light blue) and 0 (dark blue), representing the real and reconstructed data are identical and completely different, respectively. The plot shows a clear gap in the Wasserstein distance near 1, above which the KSComplement of genes, describing the reconstruction quality, is near zero. Those genes that were poorly reconstructed (109 out of 8,516, or 1.28%) were removed for downstream analyses.

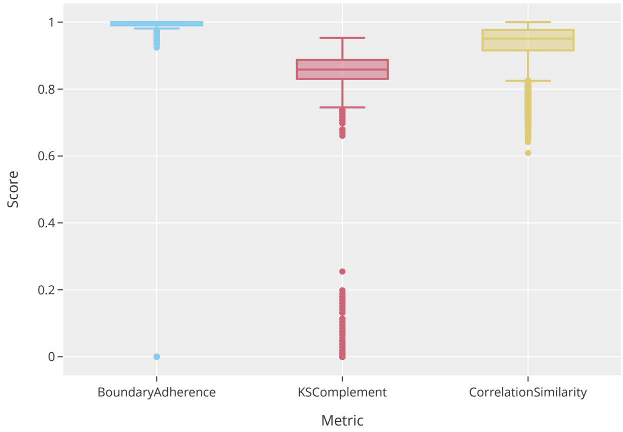


**Supplementary Figure S10**: **Assessing synthetic data with sdmetrics.** The figure shows three metrics from the *sdmetrics* Python package: Boundary Adherence, KS Complement, and Correlation Similarity. We show a subsample of one million points for the Correlation Similarity metric, due to its high computational cost. We compared the test set of real patients with the test set of reconstructed patients using our synthetic-data generation pipeline. See Supplementary Table S5 for summary statistics of each metric.

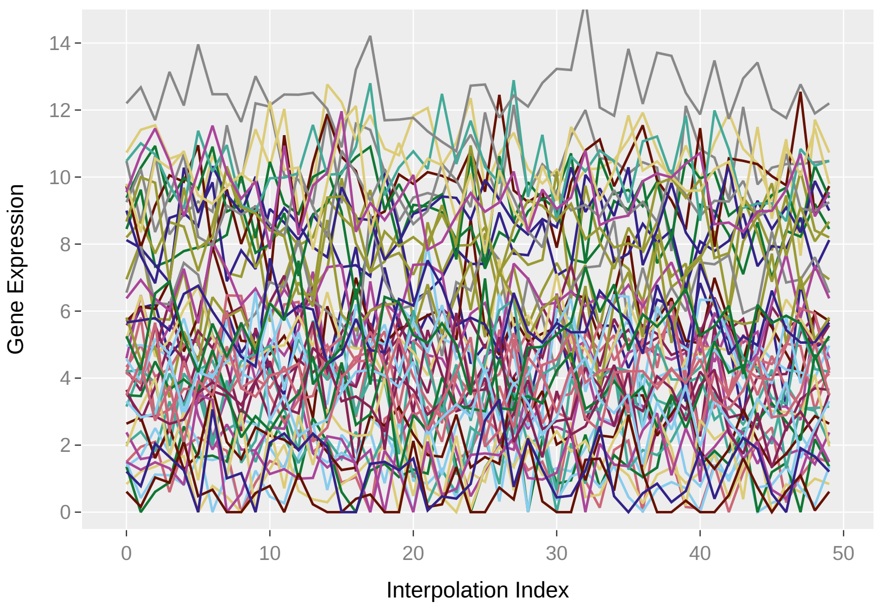


**Supplementary Figure S11**: **Control trajectory example.** Early to early stage trajectory

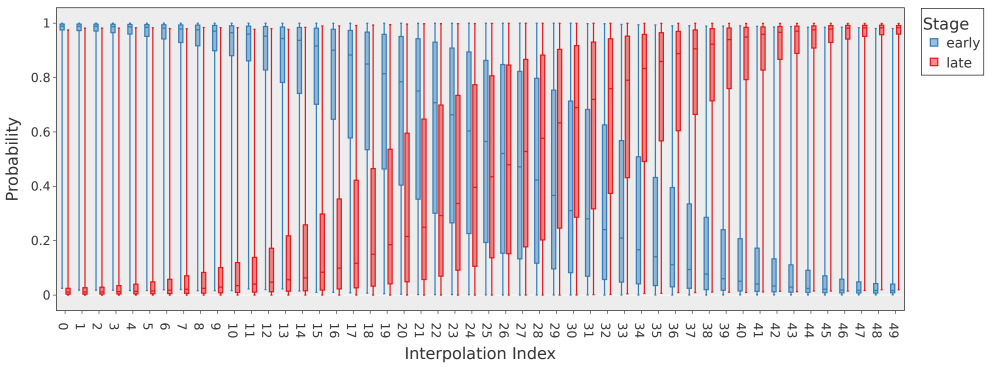


**Supplementary Figure S12**: **Classification of early and late stages along synthetic trajectories between train set patients** Distribution of probabilities the XGBoost classifier on the real patients assigns to the synthetic patients at each point of the interpolated trajectories. The best-performing XGBoost classifier was chosen to get the probabilities of synthetic patients, for each trajectory.

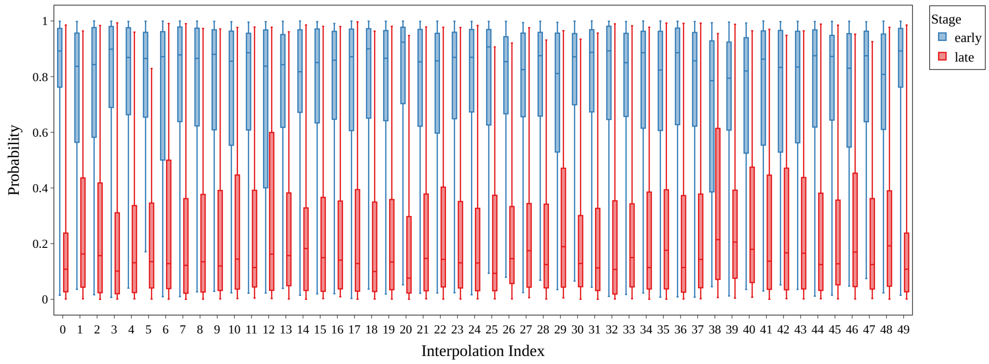


**Supplementary Figure S13**: **Control trajectory classification.** Classification of each timepoint for all control trajectories between early-stage patients.

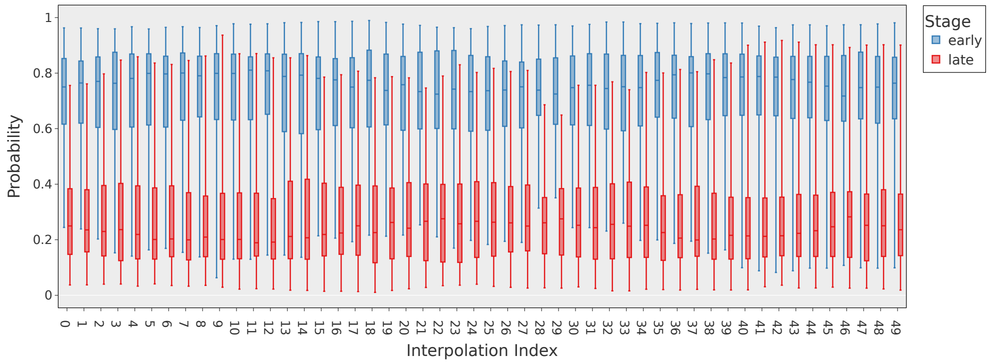


**Supplementary Figure S14**: **Classification of each trajectory point with the randomization experiment results.** The XGBoost classifier obtained in the randomization experiment (Supplementary Figure S1) was used to classify each point along all synthetic trajectories between early and late stage patients. Each boxplot shows the distribution of probabilities the classifier gives to the synthetic patients at each point in the interpolated trajectories. In contrast to the results from the non-randomized classifier (see main text Figure 3a), the likeliest classification is always early stage, which is the majority class (Supplementary Table S2).

a)

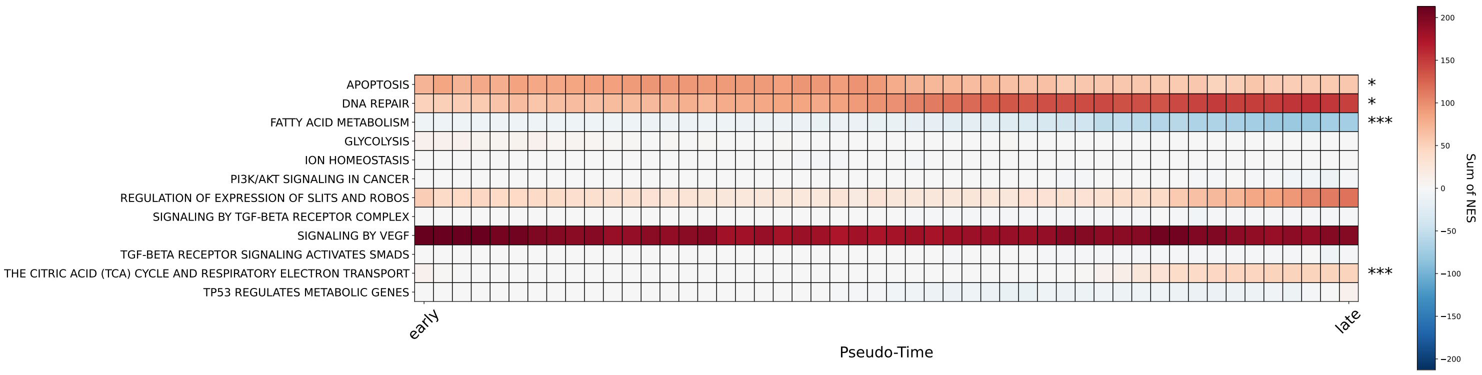


b)

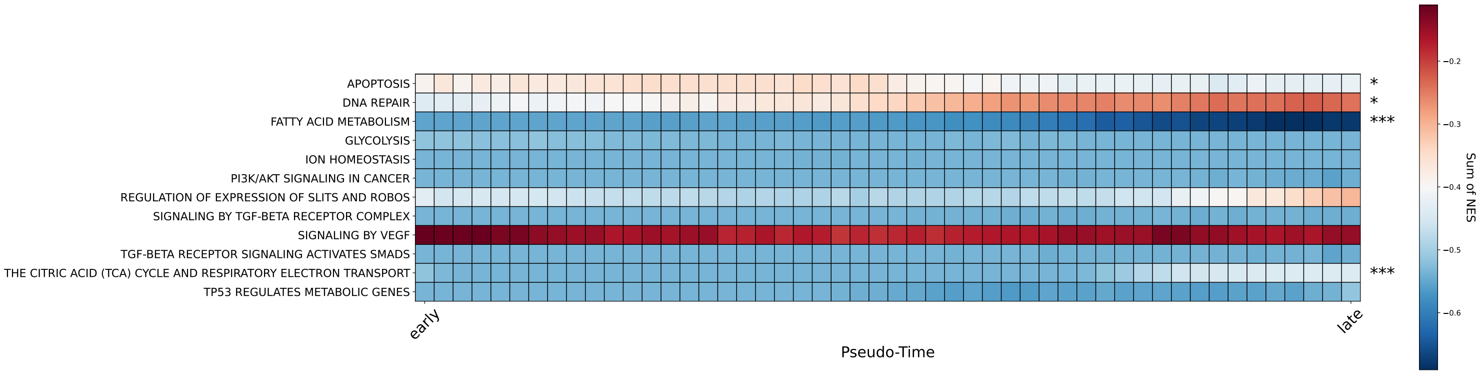


**Supplementary Figure S15**: **Relevant pathways for ccRCC found in the literature**. These are the the most relevant pathways that our enrichment procedure determined to be non-zero. Both subpanels show the same pathways, but panel **b)** has been MinMax-scaled to highlight differences in regulation along time. These pathways are related to the Warburg effect (TP53 REGULATES METABOLIC GENES and GLYCOLYSIS), PI3K, the citric acid cycle, apoptosis, angiogenesis (SIGNALING BY VEGF), among others. The equality of the distribution of the NES values between the first and last pseudotime points was tested using the Mann-Whitney U test. The p-value levels are indicated as: *, $p \leq 0.05$; **, $p \leq 0.01$; and ***, $p \leq 0.001$.

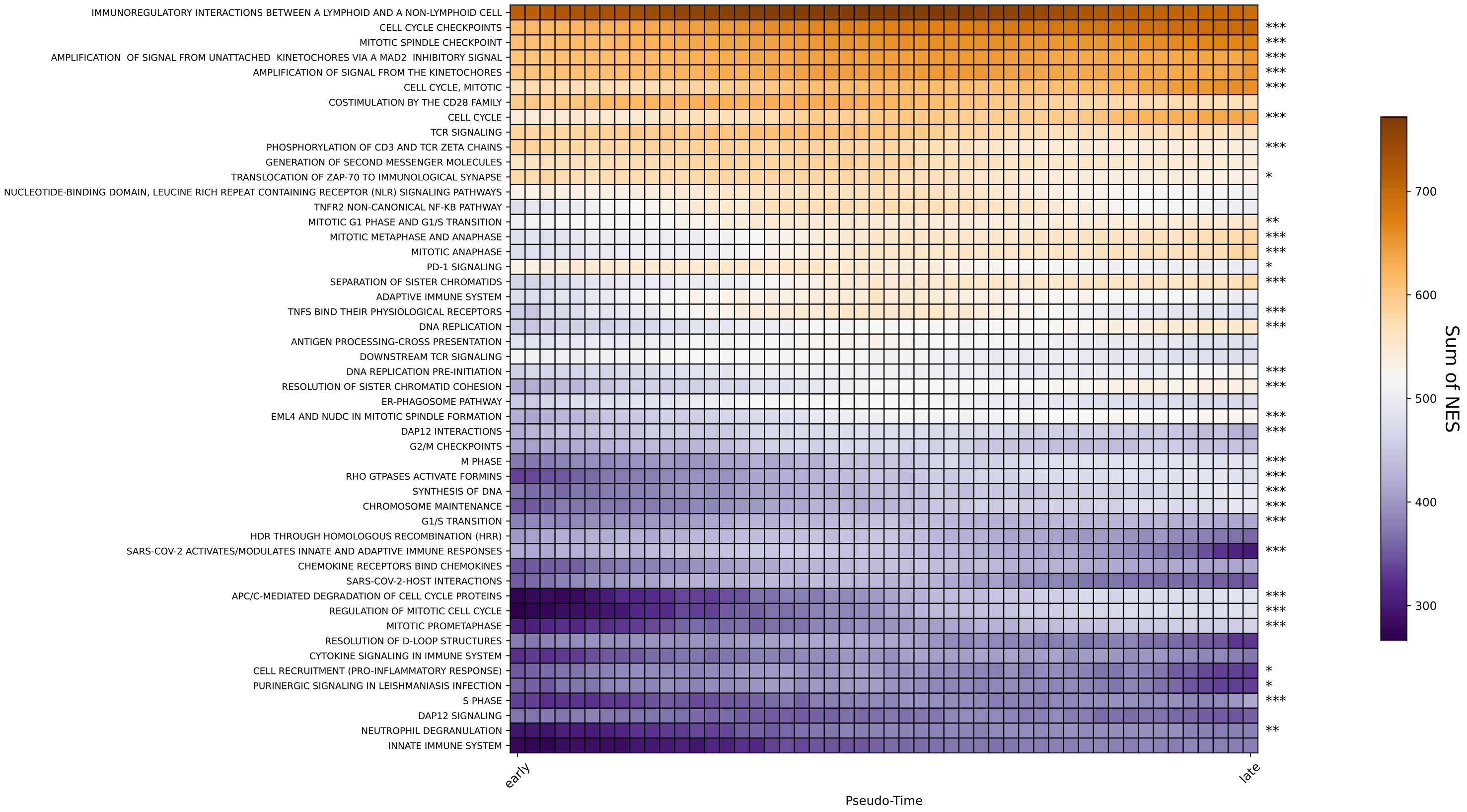


**Supplementary Figure S16**: **Top 50 most *up*regulated pathways in the synthetic trajectories**. These are (on average) the most upregulated pathways when applying GSEA with the Reactome database on the generated synthetic trajectories of ccRCC. The equality of the distribution of the NES values between the first and last pseudotime points was tested using the Mann-Whitney U test. The p-value levels are indicated as: *, $p \leq 0.05$; **, $p \leq 0.01$; and ***, $p \leq 0.001$.

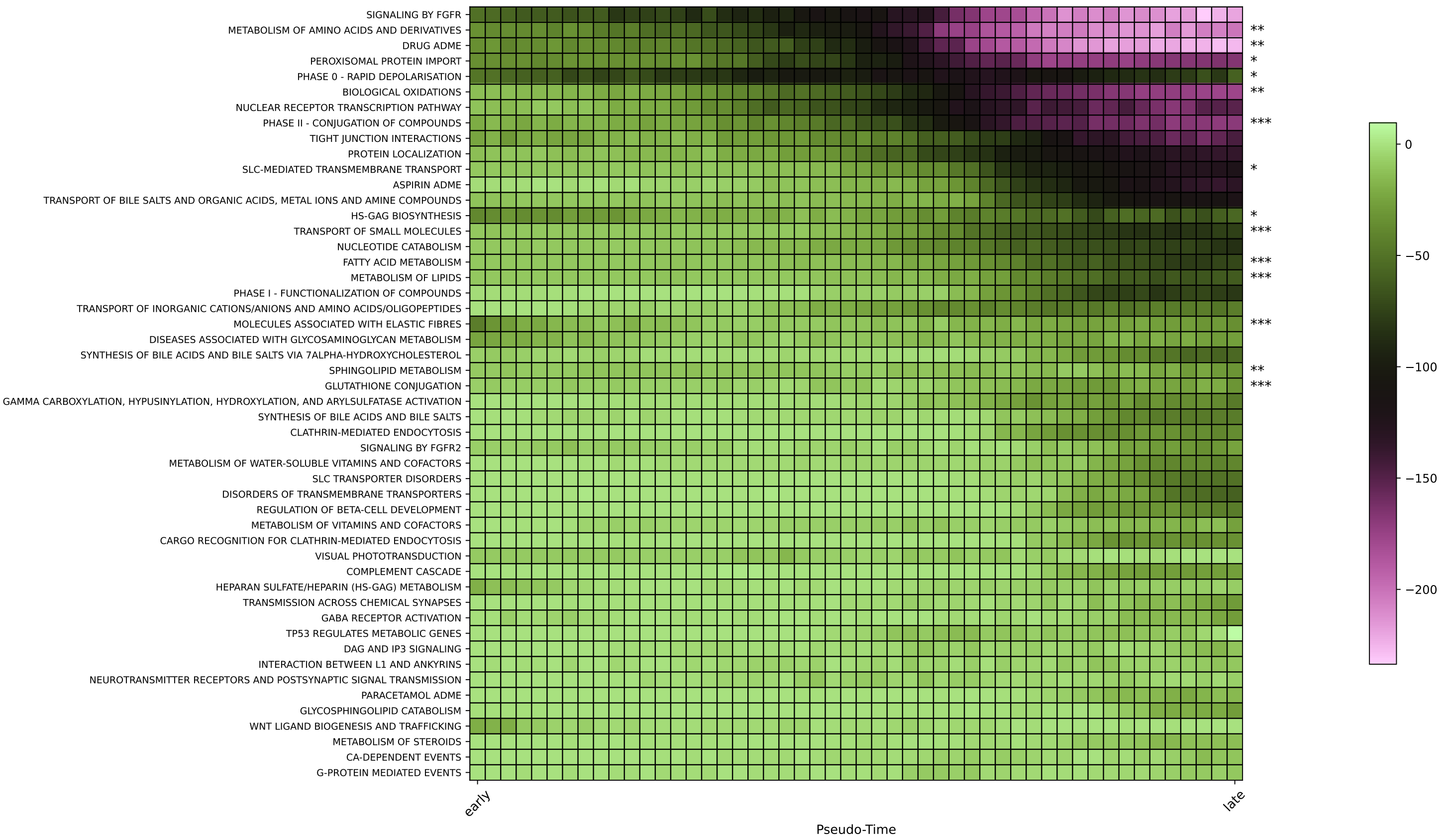


**Supplementary Figure S17**: **Top 50 most *down*regulated pathways in the synthetic trajectories**. These are (on average) the most downregulated pathways when applying GSEA with the Reactome database on the generated synthetic trajectories of ccRCC. The equality of the distribution of the NES values between the first and last pseudotime points was tested using the Mann-Whitney U test. The p-value levels are indicated as: *, $p \leq 0.05$; **, $p \leq 0.01$; and ***, $p \leq 0.001$.

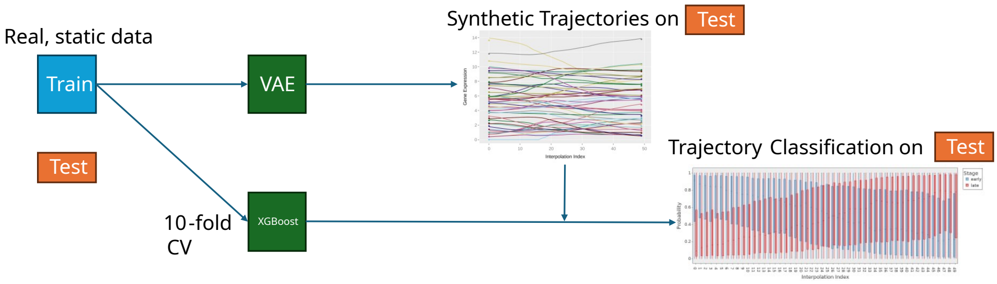


**Supplementary Figure S18**: **Strategy for avoiding data leakage in the trajectory generation and assessment.** We have split the TCGA dataset into a train (blue) and test (orange) set of patients. The same split was kept to train the VAE and the XGBoost classifier. Trajectories were then generated between patients within the set test patients, and assessed with the XGBoost classifier, which had not seen those patients before.
